# Tailored mass spectral data exploration using the specXplore interactive dashboard

**DOI:** 10.1101/2023.10.03.560677

**Authors:** Kevin Mildau, Henry Ehlers, Ian Oesterle, Manuel Pristner, Benedikt Warth, Maria Doppler, Christoph Bueschl, Juergen Zanghellini, Justin J.J van der Hooft

## Abstract

Untargeted metabolomics promises comprehensive characterization of small molecules in biological samples. However, the field is hampered by low annotation rates and abstract spectral data. Despite recent advances in computational metabolomics, manual annotations and manual confirmation of in-silico annotations remain important in the field. Here, exploratory data analysis methods for mass spectral data provide overviews, prioritization, and structural hypothesis starting points to researchers facing large quantities of spectral data. In this research, we propose a fluid means of dealing with mass spectral data using specXplore, an interactive python dashboard providing interactive and complementary visualizations facilitating mass spectral similarity matrix exploration. Specifically, specXplore provides a two dimensional t-SNE embedding as a jumping board for local connectivity exploration using complementary interactive visualizations in the form of partial network drawings, similarity heatmaps, and fragmentation overview maps. SpecXplore makes use of of state of the art ms2deepscore pairwise spectral similarities as a quantitative backbone, while allowing fast changes of threshold and connectivity limitation settings, providing flexibility in adjusting settings to suit the localized node environment being explored. We believe that specXplore can become an integral part in mass spectral data exploration efforts and assist users in the generation of structural hypotheses for compounds of interest.

**Technical Terms:**

- A network is a collection of connected features. In our case, a network consists of MS/MS spectral features connected provided their spectral similarity is high. Networks are represented using node-link-diagrams.
- Node-link diagram -a term commonly used to refer to the graphical representation of a network via nodes and links (i.e. edges). In this paper, we use node-link diagram and network-view interchangeably.
- A node is a feature in a network that can be connected to other features via edges. An alternative term for node is vertex.
- An edge is a connection between two nodes. Other terms for edges are links or vertices.
- Network layout refers to the spatial arrangement of nodes and edges on an usually two dimensional plotting surface. Network layout is also sometimes referred to as embedding. This term is avoided in this paper to avoid confusion with embedding in the machine learning sense.
- Given a network *G*(*V, E*), where *V* denotes its nodes and *E* its (weighted) edges, we define its topology as the relationships between individual (groups of) nodes and edges or the network as a whole, irrespective of the network’s layout.
- Molecular Networking (MN) is an exploratory data analysis technique merging spectral similarity-based topological clustering and visualization as node-link diagrams.
- The plain English words group/grouping are wherever appropriate to avoid jargon terms such as clustering (as in k-medoid or k-means clustering), embedding (as in projection of groups of features into a close-by lower dimensional space), or molecular families. The latter are groups of spectral data features clustered and visualized as network-views via traditional MN or feature based molecular networking (FBMN). Molecular families, usually represent smaller, disconnected networks that are part of a larger dataset. When we refer to this disconnected nature, we use the phrasing disjoint sub-network for emphasis.

## 1 Background and Motivation

Untargeted metabolomics deals with the elucidation and characterisation of small molecules in complex biological systems. Small molecules or metabolites cover an enormous chemical diversity involved in a vast range of biological functions. This chemical diversity leads to complex and heterogeneous data that is difficult to provide consistent and automated workflows for [1]. Computational metabolomics tools which assist manual data evaluation and annotations efforts such as experimental networking thus remain critical to the field [2]. Molecular Networking (MN) hosted on the Global Natural Products Social Molecular Networking (GNPS) servers is possibly the most used computational metabolomics tool for exploratory data analysis work using LC-MS/MS (Liquid Chromatography Tandem Mass Spectrometry) data [3–6]. The core idea behind MN is that, since similar structures tend to fragment similarly, spectral similarity may be used to construct spectral feature groups with implied structural similarity. In MN, the modified cosine score similarity matrix of the measured spectra forms the basis for constructing such groups using a network topology approach [3]. Nodes in the network represent MS/MS spectra features that may be connected via edges as a function of pairwise spectral similarity. Indeed, pairwise similarity thresholds and other network processing parameters are used to filter the complete network of all possible pairwise connections such that only edges for high pairwise spectral similarities remain. Using this approach, interrelated spectra form small, separated (disjoint) groups of nodes of high intra-spectral similarity commonly referred to as molecular families [3]. MN can thus be viewed as an exploratory analysis framework merging topological grouping and network visualization. Molecular families serve two separate functions, a) they provided an ordered data overview, and b) they may be used to assisting network annotation propagation, that is, the propagation of structural hypotheses from known structures to unknown ones via proximity in the network [5, 7–9],.

While the MN workflow is hugely successful [5], it comes with its own trade-offs. For instance, the use of the modified cosine score and minimum fragment overlap requirements poses a stringent similarity criterion for connectivity, resulting in rather sparse networks suitable for representation as disjoint sub-networks. However, the modified cosine score may miss structurally related analogs which exhibit larger fragmentation differences, while the disjointness of the molecular families may obscure relationships between them. Importantly, MN operates using a single global threshold setting on the basis of which connectivity may or may not exist. Such a global threshold is unlikely to work well for all chemical families measured, where some may exhibit much richer or much sparser fragmentation or overlap thereof. In addition, the combination of disjointness of spectral groupings, as well as the disconnected runs with new randomly generated layouts for each molecular family make setting comparisons an arduous task.

In this paper, we introduce specXplore, an interactive Python dashboard aimed at facilitating spectral data exploration in a flexible and local network topology tailored fashion. Unlike traditional molecular networking, specXplore was created with adjustable settings for heterogeneous and dense network data in mind. SpecXplore provides complementary and interactive visualizations that allow users to explore connections between spectral features using interactively adjustable network settings. SpecXplore consists of an importing module providing data integration capacities and a dashboard-module for interactive exploratory data analysis. The dashboard makes use of a two-dimensional t-SNE (t-distributed stochastic neighbor embedding) overview network of the full pairwise similarity matrix as a jumping board for localized data exploration. Rather than being based on modified cosine scores however [3, 5], specXplore is based on ms2deepscore pairwise similarities which can more accurately represent structural similarities between compounds based on their spectra via a deep learning based embedding representations [10]. Ms2deepscore pairwise similarities allow for structural similarity prediction on the basis of spectral data, in principle allowing grouping of similar compounds even if their spectra are dissimilar [10]. However, ms2deepscore may also introduce a much denser topology. At many reasonable threshold levels, node-link diagram representations of these matrices tend to become unreadable for the network as a whole [2]. To facilitate effective exploration of local neighborhoods, specXplore provides various interactive visualizations providing views of connectivity surrounding a feature or feature group of interest. Here, partial network drawings and matrix representations play an important role. Localized explorations are combined with the ability to quickly change thresholds to regenerate local views under the new constraints, allowing the careful expansion of neighborhood size for some node of interest. In addition to network based representations of local connectivity, specXplore provides means for a) investigating the raw pairwise similarity matrix directly, b) to investigate fragmentation overlaps across multiple spectra, and c) to inspect any joined-in metadata or chemical classifications from within the dashboard.

We will first outline the core components of the tool, their intended usage, and their rationale. This will be followed by illustrative examples on real data and a discussion the tool in the broader contexts of mass spectral exploratory data analyses.

## 2 Materials & methods

The specXplore workflow is divided into two components in the form of a data importing pipeline and visual analysis using the dashboard. The importing part provides data integration and pre-processing functionalities, while the dash part provides the interactive user interface for data exploration. The tool is available on github under MIT licence as a python package that can be downloaded and installed for local use https://github.com/kevinmildau/specXplore. We will briefly outline the dashboard’s importing pipeline and core visual components.

### 2.1 Data importing and pre-processing

Before data can be opened in the specXplore dashboard it needs to be processed using the workflow summarized in Figure 1 A. The input data for specXplore spectral data exploration are MS/MS spectral data. Basic LC-MS/MS data pre-processing (i.e., feature detection, MS/MS spectral exporting, etc.) is assumed to have been done elsewhere, e.g. [11], in order to reduce dataset size and feature redundancy. MS/MS feature data are assumed to be in .mgf (mascot generic format) format, where each entry should have a unique feature identifier, a precursor mass to charge ratio, and spectral data in the form of one or more mass to charge ratio and intensity value tuples. Spectral data are imported into Python using matchms [12] and specXplore data processing is performed (see SI). Spectral data can then be used to initialize a template specXplore object that automatically computes pairwise spectral similarities using three similarity scores: ms2deepscore [10], modified cosine scores [3, 12], and spec2vec scores [13]. The primary similarity score used in data exploration is ms2deepscore [10], which computes pairwise spectral similarities by transforming any two input spectra into the models’ deep-learning-based embedding representation. This step involves binning of spectra and recasting of binned spectra in the embedding space. Embedded spectra are compared against each other using dot product similarities. Importantly, the model is trained to generate embeddings such that the resulting dot products of two generated embeddings resemble structural tanimoto score based similarities as closely as possible. SpecXplore makes use of the pre-trained ms2deepscore model also used in ms2query, circumventing the need for users to train and validate their own models [14]. In addition to ms2deepscore pairwise similarities, modified cosine score similarities [3] and pre-trained spec2vec emebdding based similarities [13], are computed and integrated within specXplore for comparative purposes.

**Figure 1:**
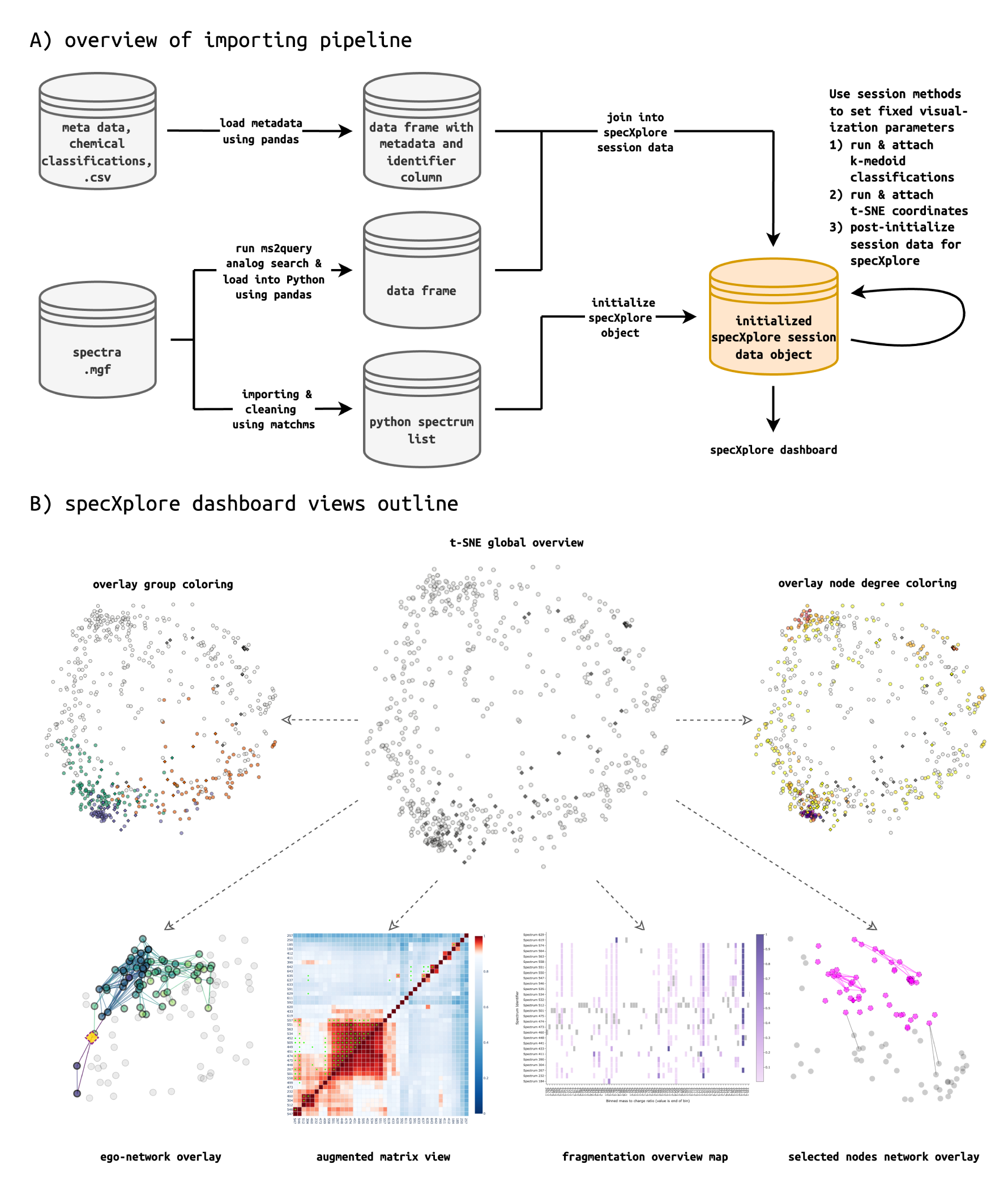
SpecXplore importing and dashboard outline. A) Flowchart of specXplore importing process. Spectra and metadata are loaded into Python, where a specXplore session object is initialized. These are loaded into Python using matchms for basic processing and used to perform basic initialization of a specXplore session data object. Here, pairwise similarities are computed for the provided spectral dataset. Optionally, metadata, chemical classifications, and ms2query library search results can be integrated into the session data object via respective join functions. Note that this requires overlapping feature identifiers across the different data frames. Before the session data object can be used, k-medoid classifications and t-SNE coordinate system setting must be added using member methods. Finally, once all processing and data joining is done, the network and plot data are initialized. B) SpecXplore dashboard overview of the main panels using the wheat data (see results section). At the center of specXplore is the t-SNE overview figure with experimental features illustrated as circular nodes and known features highlighted as darker diamonds. From this overview multiple different views portraying different detailed data views can be triggered interactively, either superimposing information onto the t-SNE figure or prompting additional figures. On the top left, nodes are colored according to grouping information, be it k-medoid or predicted chemical class. On the top right, node degrees are superimposed onto the t-SNE figure via a degree color gradient. This view grants uncluttered overview insights into topology. Starting with the leftmost bottom view, the superimposed ego-network representation for a single feature is displayed. Next to this, the Augmap displays the pairwise similarity matrix for a selection of nodes as a heatmap. Continuing, the fragmap view is shown. This view gives an overview of the fragmentation overlaps across a selection of related nodes. Finally, in the last view the superimposed network view for a larger selection of nodes is displayed. Connections within the selection are highlighted in magenta and connections out of the selection are highlighted in black. Each view is prompted interactively via node selections and button triggers, where at each view request, any modifications to the settings in the tool will immediately take effect.

The central overview of specXplore is based on a t-SNE embedding of the ms2deepscore pairwise similarity matrix. The primary tuning parameter for t-SNE embeddings is perplexity, roughly describing the average size of neighborhood expected in the high dimensional space to be considered in embedding algorithm [15, 16]. Perplexity is difficult to determine a-priori or formally tune for, hence specXplore provides functions to run t-SNE embedding routines for a range of perplexity values (usually between 5 and 50 [15]), leaving the user to select a suitable value. Perplexity value selection is assisted via original and embedded distance correlation summaries. However, the user will have to balance high dimensional distance preservation against qualitative grouping clarity to choose the ultimately used perplexity value.

While the t-SNE embedding algorithm is geared towards preserving local neighborhoods of a higher dimensional space, it does not tend to preserve global similarities well, with euclidean distances between clusters being much more difficult to interpret [17]. The embeddings thus provide the user with an abstract and condensed representation of the pairwise similarity matrix lacking clear indications of which nodes are to be considered close, and which are to be considered far away from one another. To alleviate this problem, complementary k-medoid clustering is provided using the kmedoids Python package, where clustering at various values of k provide quick glances at local neighborhoods in t-SNE overview [18]. k-medoid clustering is closely related to k-means clustering, where k-medoid omits the necessity of computing some form of centroid against which to measure distances, instead making use of the median distanced observation within a cluster as the reference. k-medoid clustering thus has a number of advantages within specXplore: a) any arbitrary distance measure may be used with k-medoid clustering, including ms2deepscore itself, b) making use of medoids circumvents the need of defining centroids, as well as any associated needs for recomputing distances from the latter making it computationally cheap, and c) since k-medoid operates directly on the similarity matrix its cluster assignments are unaffected by t-SNE projection artefacts, making it a means of data subdivision independent of t-SNE.

An optional part of specXplore is to run ms2query to provide analog library matches for all experimental spectra. ms2query makes use of a local library against which experimental spectra are compared using optimized machine learning-based matching. ms2query putative structure annotations serve as additional metadata available for each spectrum within the specXplore dashboard. Putative structures, but crucially also the chemical ontology corresponding to the determined analog can give indications about the nature of the compound represented by the spectrum in question. Taking this approach further, the analog classifications may be supplied to specXplore’s classification table, providing a rough, indirect analog based classification for all experimental spectra that can be used to identify chemical regions of interest. Of course, other tools for library matching or more direct class predictions using tools such as CANOPUS may also be used and integrated into the metadata table and classification tables of specXplore [19, 20]. Integration of these tools’ class predictions can be achieved easily using specXplore’s provided a) the chemical classification is available in tabular form and b) feature identifiers matching those supplied with the mgf file are available for each classification.

### 2.2 SpecXplore interactive dashboard

SpecXplore provides the user with a variety of views and interactive navigation options (Figure 1 B) [21]. We envision specXplore’s functionalities can be used in an interactive local exploratory fashion, where settings and their impact on local neighborhood can be evaluated seamlessly. This local exploration starts at a two-dimensional t-SNE overview figure rendered using dash and dash-cytoscape [22–24]. Here, the t-SNE algorithm is used to generate a two-dimensional representation of the ms2deepscore pairwise similarity matrix such that local neighborhoods in high dimensional space are preserved in the two dimensional plane (Figure 1 B). The two-dimensional embedding overview visualizes all points as nodes but omits any edges between nodes. Node specific feature information can be obtained via mouse-hover operations. Any features with known structures and in-silico spike in reference standards are visually highlighted via shape, color, and opacity, providing immediate pointers to areas of interest in the two dimensional projection. In addition, chemical classification and clustering information is available for visualizing up to eight different chemical or k-medoid classes at a time, where eight is set to prevent colors from becoming difficult to distinguish [25, 26]. In following Shneiderman [27] mantra of “Overview first, details on demand”, specXplore allows for the selection of any node, or collection of nodes, for more in-depth analysis in complementary visualization approaches falling into two broad categories: a) topology views, and b) data details views.

Four different topology based views are provided. First, since every node in the overview projection also exists in a network context, a node degree visualization was added. This visualization portrays the degree of each node using a color scale ranging from the lowest non-zero degree to the highest degree within the overview plot. This provides the user with immediate and global insights into the number of edges connecting to each node using current network construction parameters. This Node degree visualization, as well as edge weight distribution plots included in the settings panel, follow the principle idea of Willet et al.’s *Scented Widgets* [28], providing intuitions about the impact of settings on topology (see SI).

Second, a heatmap portraying a slice of the ms2deepscore pairwise similarity matrix for a selection of nodes is provided. Here, ms2deepscore scores are quantitatively portrayed using a divergent color scale around the current threshold setting, while exact numeric values are available via mouse-hover panels. This view is called augmap (AUGmented HeatMAP) in specXplore since it also incorporates implied adjacency matrices for modified cosine score and spec2vec score matrices at current threshold settings via additional markers and mouse-hover information, providing an effective means of comparing local neighborhood topology across similarity measures from the ms2deepscore anchor point. SpecXplore further allows for network edges to be visualized on top of the t-SNE two-dimensional layout via interactive selection and prompting.

As the third type of visualization specXplore thus makes use of superimposed ego-networks; visualizations suitable for studying a network’s topology relative to a single node [29, 30]. SpecXplore’s egonet representation allows for the visualization of all edges connecting to a single selected node at current edge threshold limitations. The visualization can be further extended by increasing hop-distance allowances, where at current threshold constraints, all edges connecting to the current node, but also all edges connected to the connected nodes, and so on, are highlighted. This allows the user to explore a branching view of network connectivity emanating from the ego node. Egonets are aimed at allowing the user to find nodes connected to a reference standard, to find neighbors of a node of potential interest, as well as to evaluate inter-group connectivity when using higher hop distance allowance usually resulting in radiating branch connections through related parts of the t-SNE overview.

Fourth, a multi-node selection based network view for intra-group connectivity assessments highlighting all edges within a selection of nodes, as well as those connecting outwards of the selection, is available for quick assessments of topology in entire areas of the t-SNE overview.

In addition to the topology views, spectral data detail panels are provided. For any group of spectra, metadata information can be presented as a table, and MS/MS spectrum plots can be generated. For pairwise spectral comparison, so-called mirror plots depicting one spectrum on the positive y axis and the other on the negative y axis are available. However, the latter are not suitable for multi-spectrum comparison and evaluations of fragmentation overlaps across more than two spectra. To support users with multi-spectrum comparisons we’ve implemented fragmentation overview heatmaps (fragmap). To generate a fragmap, we sort binned mass-to-charge ratios in ascending order and factorize for them for the x-axis, which allows to a) reduce the amount of unused white space in each individual spectrum plot by putting ascending fragments immediately next to each other regardless of mass differences, and b) to separate crowded areas of the mass-to-charge ratio axis into more easily separable pieces. The y-axis in fragmap is used for aligning the different spectra, while a color gradient is used to highlight fragment intensity. Lower intensity fragments are as visible and as straightforward to check for alignment as large intensity fragments. In addition, neutral losses are indicated as constant colored blocks inside the same visualization, providing a rich view of the overlaps in fragmentation patterns. Fragmap thus provides immediate insights into spectral overlaps in a single, concise overview. Naturally, fragmap also highlights lacks of overlap via empty x-axis lanes for spectra lacking overlap. Within the local exploration framework of specXplore, fragmaps facilitate quick assessments of spectral overlaps for local connectivity.

## 3 Results

We illustrate specXplore by applying it on real LC-MS/MS untargeted metabolomics data from two experiments: a) wheat plants LC-MS/MS data [31, 32], and b) urine metabolome LC-MS/MS from a polyphenol exposome study [33]. In both cases the approach and illustrations are similar. We’re hence focusing on the wheat data here and refer to the supplement for the briefer urine data example. Spectral data from .mgf files was loaded into the pre-processing jupyter notebooks and processed into a specXplore session object https://github.com/kevinmildau/specxplore-illustrative-examples. Upon opening the app and loading the data, the user faces an interactive two-dimensional projection of all spectra created by t-SNE. Each spectral feature in the dataset is represented as either as a circular node or highlighted as darkened diamonds for reference standards in this example. On its own, the t-SNE overview figure is difficult to read. Only a limited view of clustering trends can be observed through denser node regions and positioning of reference standards (Figure 1 B central view). This being the case, the t-SNE overview figure still provides an excellent basis for in-depth exploration of the data via interactivity. The most useful top-level exploration features are k-medoid clustering at low values of k (Figure 3 A), chemical classification (Figure 3 B), and node degree visualizations (Figure 2 A).

**Figure 2:**
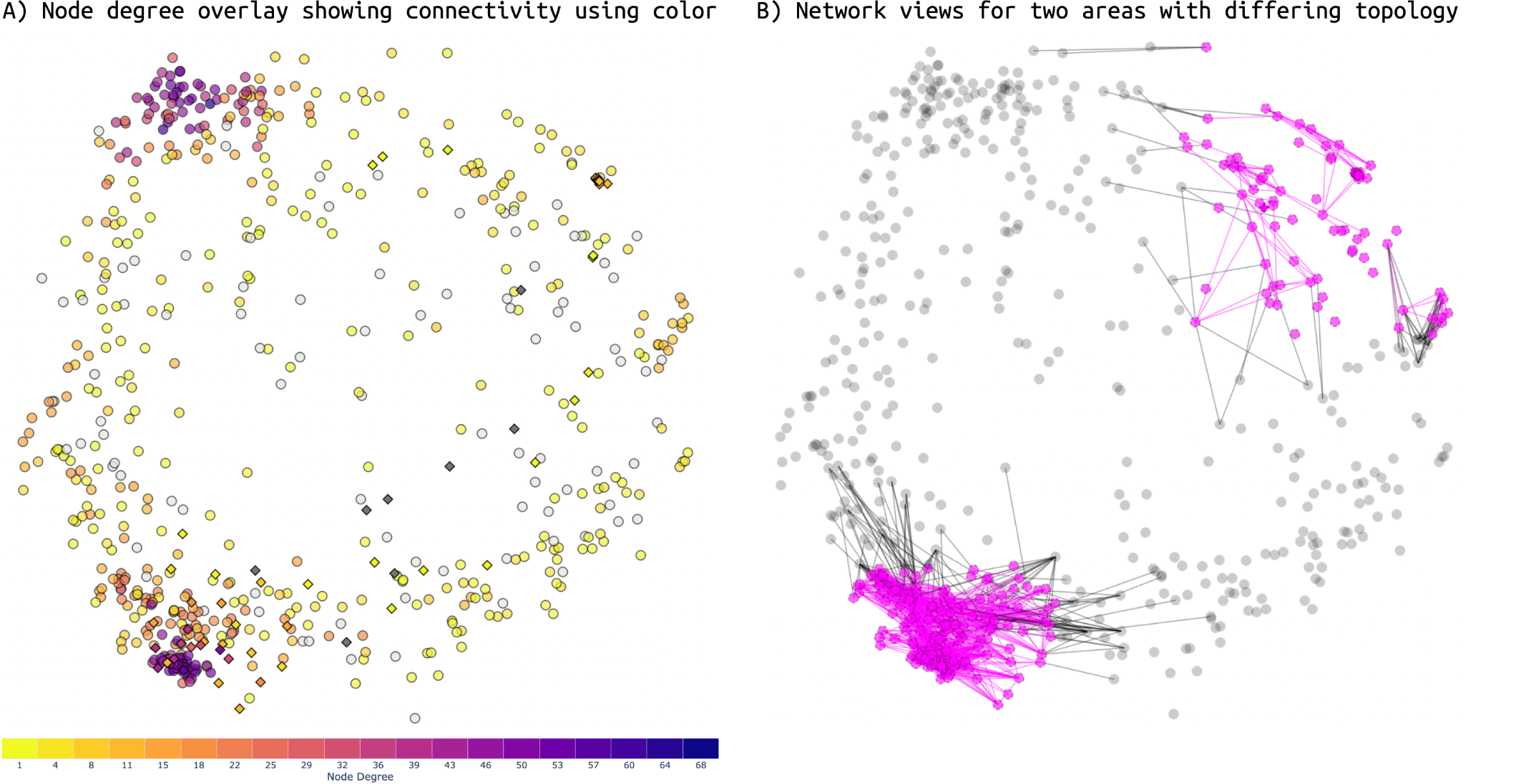
A) Node degree visualization for threshold settings of 0.8. Color indicates node degree in qualitative brackets of 10, going from lowest node degree to the highest. At this threshold, the highest node degree is 68 while the lowest visually highlighted degree is 1. Node degrees of zero are off color scale via grey color. Regions with high network connectivity are clearly visible, as are changes in topology in response to threshold changes. B) At identical threshold levels different parts of the data exhibit vastly different connectivity. For example, at threshold levels of 0.8 the lower left corner of the two-dimensional embedding is densely interconnected, while the top right corner shows sparse connectivity.

**Figure 3:**
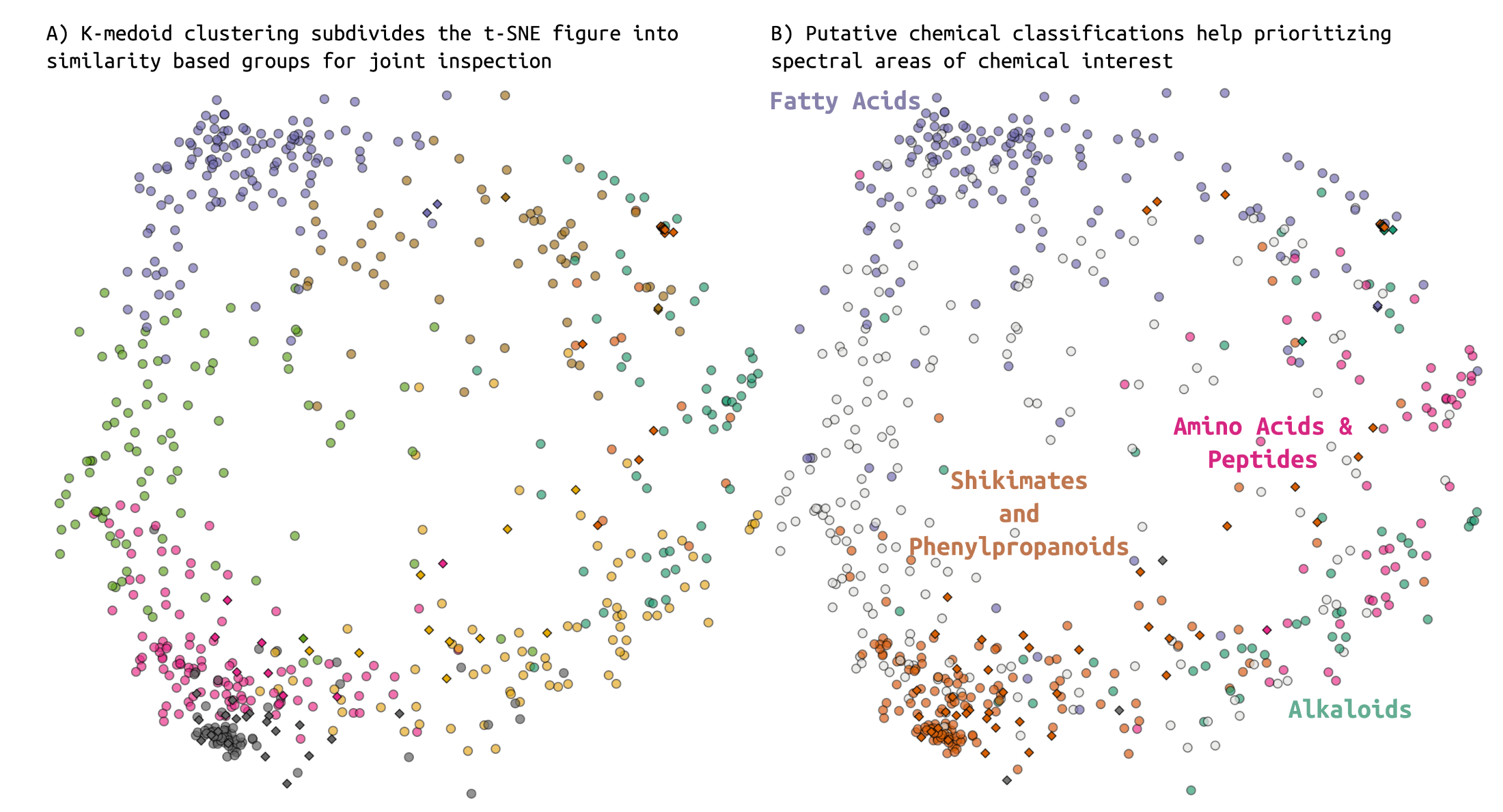
SpecXplore t-SNE overview figure with superimposed classification informaiton. A) K-medoid cluster numbers zero to seven of a clustering with k set to eight are all highlighted in color, exhausting the provided color scale. Clusters provide complementary insights into what can be considered roughly proximal in the t-SNE projection. B) Making use of ms2query analog classifications as proxies for chemical classification we can see that analog matches for our spectra already show consistent chemical organization in the data (using NPClassifier pathway ontology).

Node degree visualization provides an immediate and uncluttered view of network topology at current threshold levels, providing insights into topological groupings in the data (Figure 2 A). In addition, node degree visualizations provide the most straightforward assessment of the impact of threshold changes on local topology, while avoiding the computational cost and visual clutter of potentially large numbers of edges (see also Figure 2 A and B).

Color based highlighting of groupings based on k-medoid clustering or chemical ontology predictions can provide additional means of determining local areas of interest in the t-SNE overview. Here, k-medoid clustering provides an edge-threshold independent means of dividing the feature space into smaller groups while still making use of the pairwise similarity matrix. K-medoid clustering tends to show good agreement with both t-SNE projections and topological insights, while it provides a means of determining neighbor sets of nodes of interest (Figure 3 A). At low k values k-medoid clustering tends to produce larger data groupings (Figure 3 A), higher values of k tend to subdivide the data into smaller, localized clusters often corresponding well to topological connectivity at higher thresholds (see supplementary information). Putative chemical classifications may serve a similar means of prioritizing areas of interest in the t-SNE embedding.

In addition to the use of various color highlighting approaches to detect groupings of interest or achieve a birds-eye view of topology, specXplore makes use of edge overlay visualizations granting insights into local node connectivity patterns at adjustable similarity threshold settings (e.g. Figure 2 B). These views provide insights what nodes are considered adjacent to one another given current similarity thresholds. While network views provide import simplified views of the similarity relationship between nodes, augmap views provide deeper insights into the similarity matrix and quantitative backbone of all of specXplore’s visualizations (Figure 4). These quantitative insights can be used to adjust local threshold settings accordingly, or be used as an alternative to network views for small node selections altogether.

**Figure 4:**
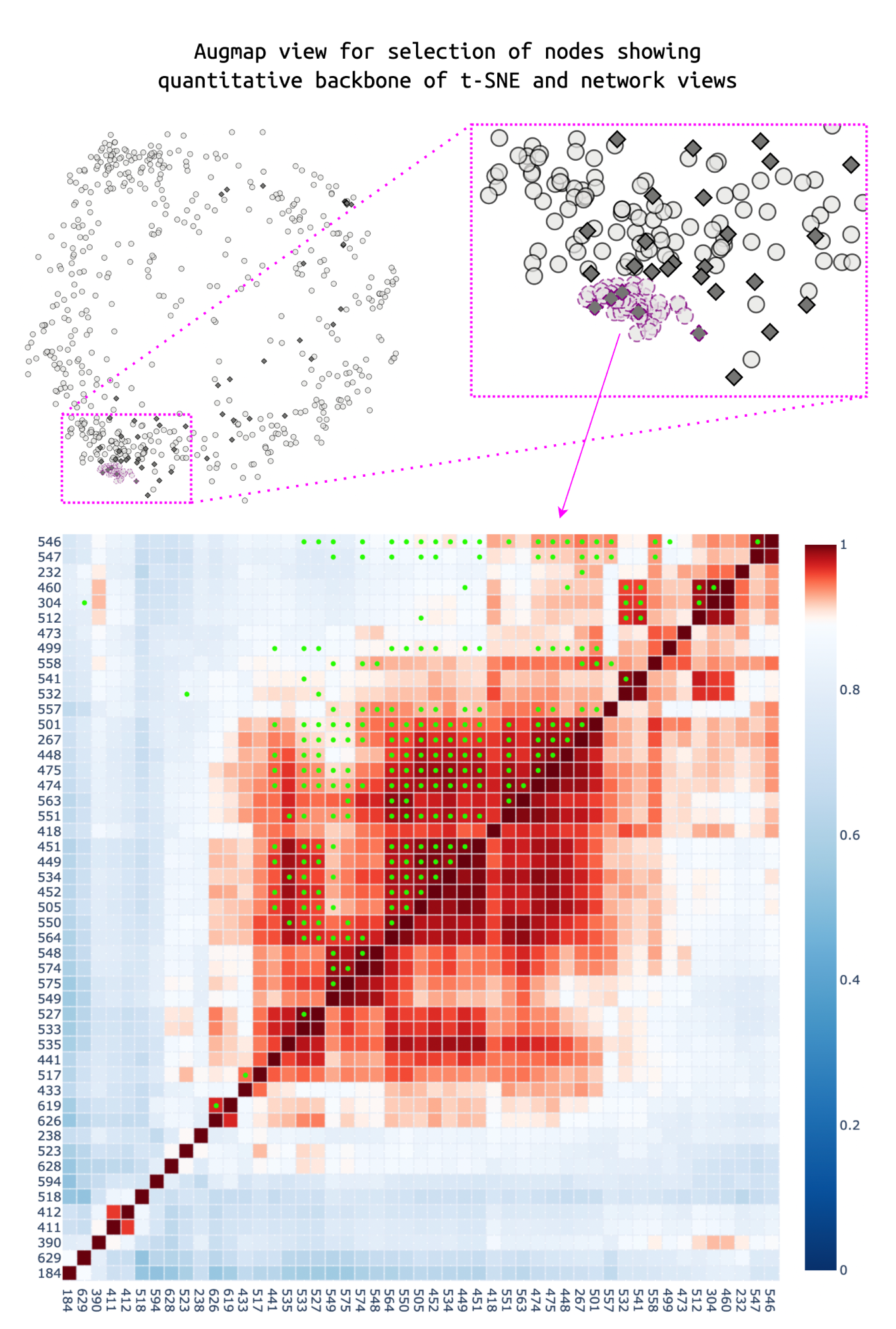
Pairwise spectral similarity and score adjacency matrix agreement visualized as marker augmented heatmaps. Feature identifiers are repeated on x and y axis, while the color gradient indicates quantitative similarity for the primary score, i.e. ms2deepscore. Here, a small selection of nodes in the bottom left area was selected. At threshold level of 0.9 for ms2deepscores, many of the selected nodes are connected via edges, indicated as red in the divergent color scale around the threshold. The remainder of the nodes tend towards light blue and white color, indicating that lowering threshold to 0.8 or 0.7 would lead to almost all pairwise connections being above threshold. In addition, we can see that at current numeric threshold value of 0.9, adjacency matrix (i.e. which pairwise relationships form edges or not) disagreement is high via the lack of or presence of circular (modified cosine score) and rectangular (spec2vec) markers. In this area of the graph, ms2deepscore appears to find the strongest connectivity patterns, while its adjacency matrix is missing some of the cosine score connections at threshold 0.9. No spec2vec connections are present at current thresholds, indicating that the pre-trained model covers the selected node subset poorly or that the score requires lower thresholds for connectivity to become apparent. Hover pop-ups are available to assess exact scores across the three scoring approaches for quantitative insight.

While specXplore’s global overviews provide insight into rough patterns of the data, its localized views provide more detailed insights into possible relationships between features in the wheat data. There are numerous ways to delve further into the data given areas of interest have been found. One sensible approach in specXplore is to explore connectivity around known reference standards. For instance, exploring the densely connected feature area around the Procyanidin-A2 corresponding feature (Figure 5 A), we can see characteristic fragment ion overlap as well as much higher complexity of the experimental spectra (Figure 5 B & C). Such overlaps in fragmentation patterns and implied substructure overlaps alongside local topology may and other metadata information may serve as vital starting points for MS/MS structural hypothesis generation and manual annotation efforts. Which features are considered of interest, and how stringent overlaps will need to be to be useful, naturally depends on the analysis goals.

**Figure 5:**
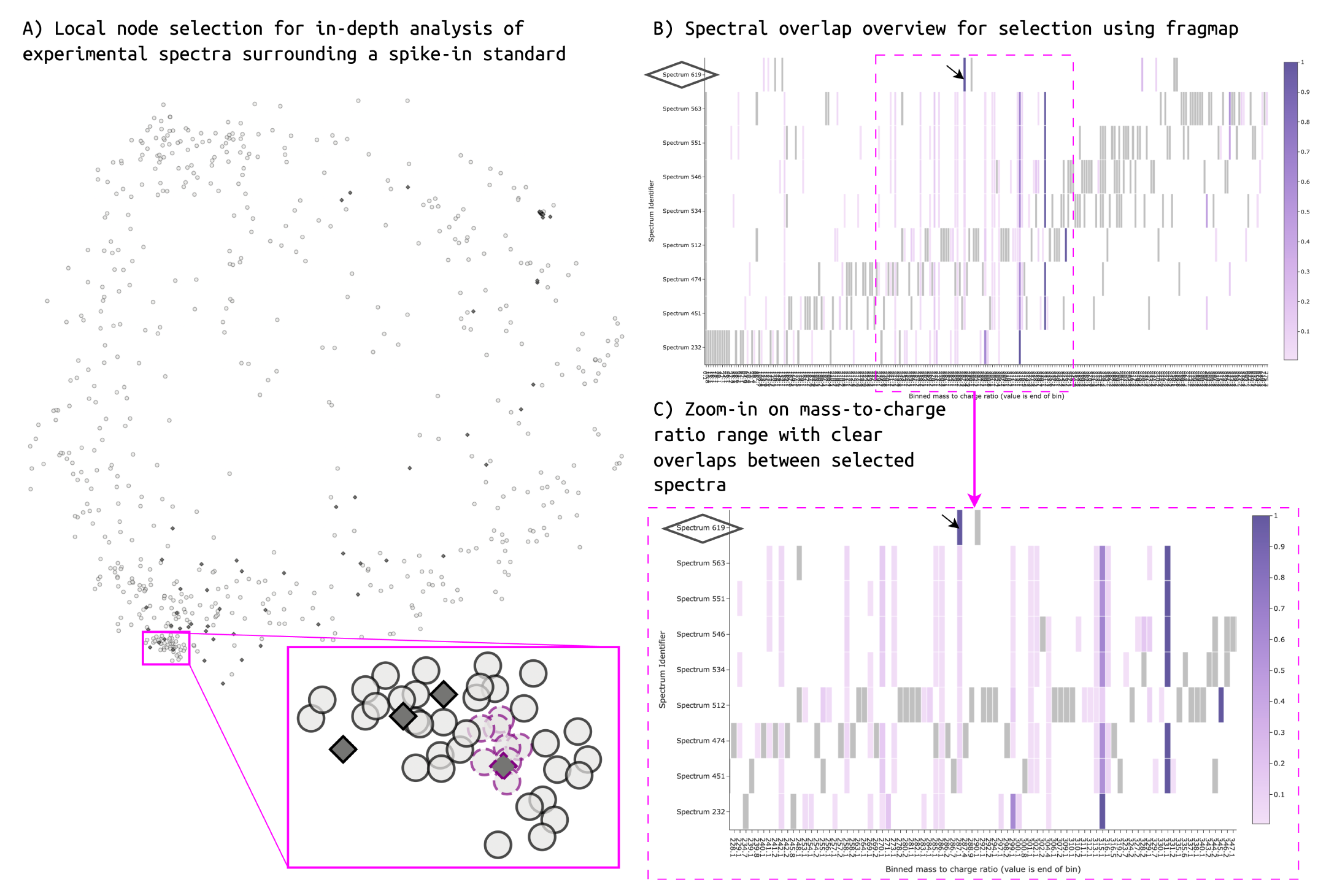
The process of exploring local environment of a known feature node is illustrated. A) Procyanidin-A2 reference standard surrounding feature selection in lower left corner of the t-SNE embedding. The selected nodes are highlighted using magenta outlines. The selected reference standard is highlighted as a dark grey diamond with dashed selection outline in magenta. B) Fragmap view of the fragmentation overlap between spectra selected in A. The reference standard spectrum is of significantly lower complexity than the experimental spectra. Only few fragments appear overlapping. However, many fragments are shared across experimental standards in a clear pattern indicating structural relationship. The reference standard is highlighted on the y-axis using an added diamond outline. The highest intensity fragment for the reference standard is further is highlighted using an arrow. C) Zoom in on overlapping area in fragmap view. All except one experimental feature in the local selection shares a low relative intensity fragment ion at mass 287.055 that appears to correspond to the most intense fragment in the reference standard.

## 4 Discussion

We’ve developed specXplore with two primary aims in mind. First, we wanted it to provide a flexible data exploration approach for structural hypothesis generation and manual network annotation propagation. Here, we specifically wanted to to avoid inflexible exploration settings and corresponding fixed groupings at odds with exploratory data analyses such as single threshold settings or the creation of a singular sub-grouping of the data. Second, we wanted specXplore to provide a means of understanding and tailoring network processing settings to the local data at hand. In keeping with these aims, we’ve developed specXplore as an interactive dashboard that supports flexible local data exploration using complementary visualization approaches and on-the-fly adjustments of settings in a well anchored context. In specXplore, no particular subdivisions are created automatically, nor are there topological parameters to ‘fix’. We opted to approach the data exploration in this way to accommodate data heterogeneity better. We think of specXplore as providing an interface to a heterogeneous and complex network of spectral similarities. This network aims to present pairwise structural similarities between features via ms2deepscore, but is also impacted by differential fragmentation behaviors for different compound classes or groups, where some may fragment richly with strongly overlapping characteristic fragment ions, while others fragment in a more limited and erratic fashion. In addition, some chemical classes may be covered better or worse by the ms2deepscore model. With data this variable, local exploration and local setting tailoring seems the most sensible. In practice, specXplore requires the user to delve into spectral similarity data of and find localized target groups based on their own criteria of interest and tolerances in spectral and implied structural similarity. While certainly requiring more effort from the user, this provides them with unprecedented flexibility.

During the development of specXplore we drew inspiration from two extensions of molecular networking (MN): MolNetEnhancer and MetGem [34–36] (see Figure 6 A-C). MolNetEnhancer extends MN by providing a data integration component facilitating the identification of molecular families of interest. Here, molecular families are color coded to the most dominant spectral motif inside the family. This effectively joins ms2lda ‘classifications’ and topology information and represents said information inside molecular networks using Cytoscape [37, 38]. MetGem on the other hand improves upon MN by providing a complementary t-SNE overview representation of the whole network based on the pairwise similarity matrix without thresholds, potentially better preserving relationships between groupings of spectra [15, 35]. Both MolNetEnhancer of MetGem have found use in the field (e.g. [39–41] and [42–44]) and highlight the potential in extending and tailoring the MN workflows. In addition, specXplore draws inspiration from the EdgeMaps network visualization approach [45, 46] (see Figure 6 D). In EdgeMaps, a complex and dense network is embedded in a two-dimensional projection of the node similarities, and directed edges between any interconnected nodes are visualized only upon interactive demand. SpecXplore makes use of the embedding approach of MetGem and EdgeMaps in order to create a meaningful node layout basis around which to explore topology dynamically, using interactive prompting as in EdgeMaps to highlight local topology. Chemical space exploration and prioritization is facilitated through chemical-classification-based node coloring as done in MolNetEnhancer, but also using visual highlighting of known features. SpecXplore further draws inspiration from Network Annotation Propagation, be it automatic or manual [8, 47, 48], where we designate the primary aim of specXplore to not be to assist in the subdivision of the data, but rather the exploration of local topology with the end goal of structural hypothesis propagation and spectral cross comparison. Combining elements of all these approaches, specXplore is a uniquely flexible data exploration approach for mass spectral data that covers a broad range of complex network visualization tasks [49].

**Figure 6:**
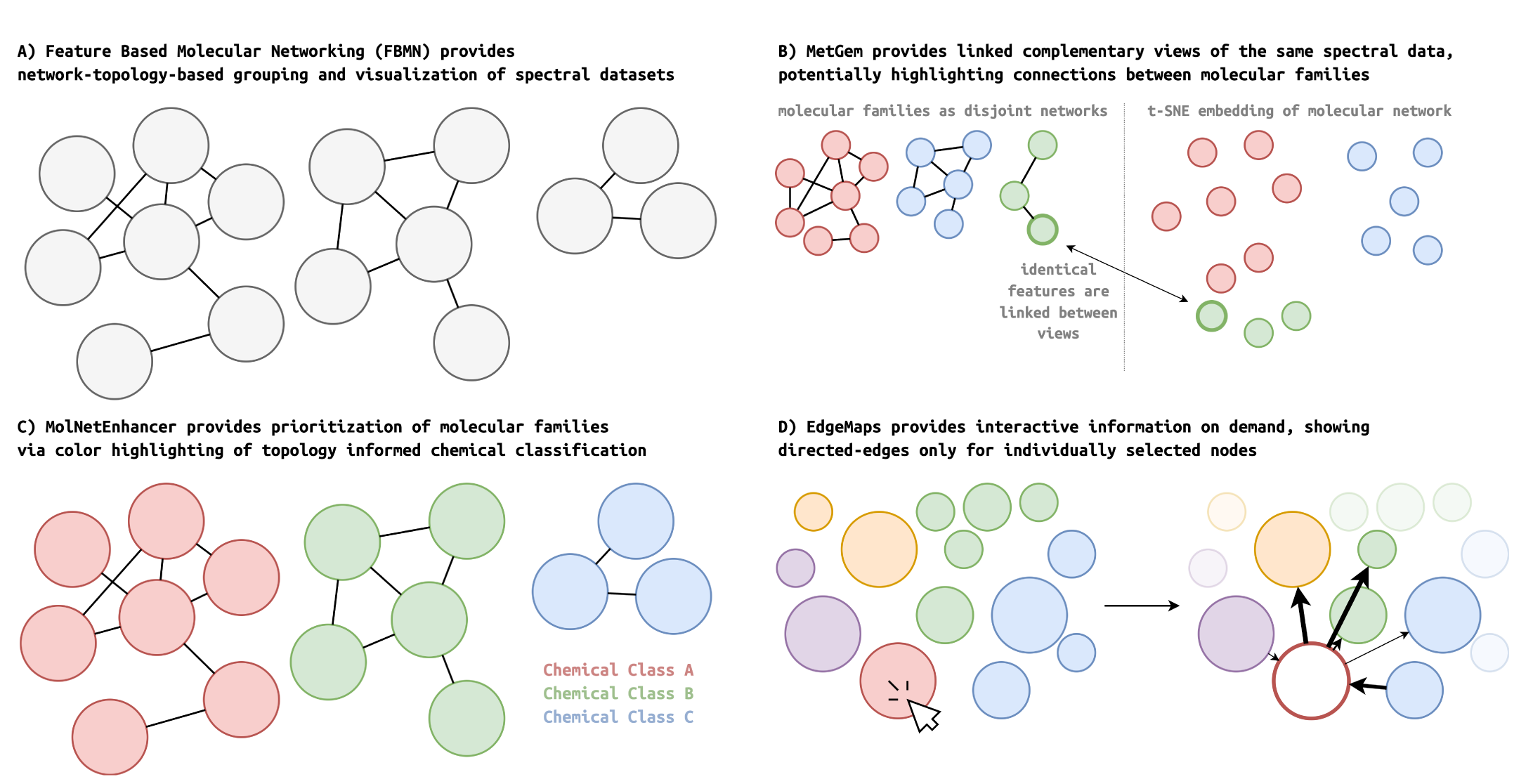
SpecXplore is a exploratory dashboard aimed at exploring MS/MS data generated from LC-MS/MS datasets. It is inspired by and related to A) FBMN (Feature Based Molecular Networking) through the premise of molecular networking [5], B) MetGem through the use of a low dimensional embedding in a complementary fashion alongside network views [35], C) MolNetEnhancer (Molecular Network Enhancer) through the idea of joining color and classifications to help users prioritize sub-networks of interest [34], and D) edgeMaps through the approach to unlocking details including edges in the network views in an interactive fashion [45]. All illustrations are conceptual simplifications and do not represent the exact visual choices of the tools in the respective papers.

### 4.1 Impact of settings in traditional molecular networking and specXplore

MN, as hosted on GNPS, provides an interesting and successful framework for topology based mass spectral exploratory data analysis (EDA) [4]. While EDA is characteristically dynamic and flexible, the settings and visual angles used can have a tremendous impact on how the data is viewed and used. The primary topological settings in MN are a) the pairwise similarity thresholds used for adjacency determination as well as corresponding minimum fragmentation overlaps, b) top-K neighbor limits on individual nodes, and c) maximal molecular family sizes. Additionally, one can consider the choice of the modified cosine score (or spec2vec scores) as the underlying metric a setting. The particular combination of default settings and similarity metric used in the GNPS workflow is used to create a subdivision of the spectral data into disjoint groupings that are visualized separately as sub-networks. Indeed, modified cosine similarity matrices will tend to be sparse, while default thresholds of 0.7 will lead to even sparser adjacency matrices. Furthermore, hub nodes connecting different groupings will be limited in scope by top-K limits and limits to group sizes. Naturally, settings for individual runs can be changed. In general however, the combination of settings and visualization approaches are tailored to accommodate sparse representations and not dense networks. A consequence of this is that molecular families represent groups of very high spectral similarity with no visible links to other families. Missing connections between nodes and clusters through restrictive settings for local topology, as well as hard to decipher hairball molecular networks through liberal settings may easily be encountered. It is hence a key issue for users of MN that GNPS reruns with different settings are slow, while comparisons of runs from one to another is non-trivial. Indeed, MN produces disjoint molecular families to be analysed separately, while reruns with different settings produce different molecular families in size, composition and natural ordering. In addition, individual molecular families make use of randomized force-directed layouts, leading to different node positioning in each run and family.

This means that the preservation of any kind of mental map of the different runs and how they compare against one another is difficult. While the speed bottleneck of traditional MN on GNPS can be partly overcome with tools such as MetGem or MZMine [11, 35], which allow fast local reruns separating data processing from networking settings, the comparison of different runs to one another still remains difficult.

Unlike traditional MN, specXplore was created with dynamic settings for heterogeneous and dense network data in mind. This is reflected in a local sub-network focus, as well as in the use of a fixed, meaningful two dimensional layout use in the form of a t-SNE projection as done in [35]. The use of fixed coordinates as layout allows the user to create a mental map of the data, as well as allows them to study the impact of changed network settings on their requested local views easily against the thus provided visual anchor-point.

The underlying pairwise similarity matrix of specXplore is provided by a ms2deepscore model [10]. This model is trained to predict pairwise structural similarity from spectral data is used to provide better capabilities for linking spectra with stronger fragmentation differences, albeit at the cost of denser networks that are more challenging to visualize. Any MN approach will be heavily impacted by the choice of spectral similarity measure, possibly leading to very different network topology and connections across different scores. While different scores are being investigated with respect to their library matching performance, there are no direct evaluations of suitability for MN like applications to our knowledge [1, 10, 50, 51]. In specXplore, ms2deepscore is used because of its in-principle capability of connecting spectra of similar metabolites even in the presence of larger fragmentation differences. However, since ms2deepscore is a machine learning model it can only be expected to work well for spectra and metabolites well covered by its training data. To accommodate this deficit at least in part, specXplore provides the augmap views granting insights into differences between ms2deepscore, modified cosine scores, and spec2vec scores on the same data.

### 4.2 Dense networks and network layout choice

SpecXplore is a tool for interactive data exploration with a pronounced network aspect. This means that the choice of positioning of nodes is in part a question of network layout choice. In network visualization literature the task of laying out dense networks of many nodes and edges is generally appreciated to be a notoriously difficult task. Commonly, force-directed layout algorithms, such as extensions of Eades [52], Fruchterman-Reingold [53], or Kamada-Kawai [54], are employed owing to their availability and computational tractability [55]. However, by not directly considering network aesthetic criteria in their cost functions, these approaches do not scale well visually to dense or large networks, producing hard-to-read, or even completely unintelligible, network renditions [55]. Many factors contribute towards this poor readability [56], most notably the number of edge bends and crucially the number of edge crossings [57, 58], even in larger figures [59]. In the worst case, naive attempts to visualize such networks as (force-directed) straight-line node-link diagrams result in so-called “hairballs”; dense, unintelligible clusters of nodes and edges, which do not allow for meaningful exploration of either local or global topology [60, 61]. In specXplore, we tackle the layout problem by combining a latent variable space embedding [35, 62] of nodes with interactively triggered partial network drawings [45, 63]. The latent variable embedding serves the purpose of a fixed jumping board for localized explorations. In addition, unlike for force-directed methods which merely focus on minimizing node-node and edge-node occlusions, as well as edge-crossings, node positioning in t-SNE is based explicitly on pairwise similarity, making node position in itself an informative quantity similar to the approach in EdgeMaps [45]. In general, a t-SNE embedding may be considered a simplified similarity matrix visualization in its own right without needing any additional network layers. The choice of using the t-SNE map to access additionally enriched views, including sub-network views for selected subsets of the complete network, is in line with Shneiderman [27]’s mantra of “*Overview first; details on demand* “. Dense hairball visualizations or edges traversing the whole t-SNE embedding are avoided by only visualizing edges on demand, as well as by allowing users to set thresholds such that local connectivity is visually manageable and scientifically useful. In addition, settings to reduce visual clutter similar to those used in the GNPS version of MN exist to reduce the potential for overbearing numbers of edges. For instance, top-K limits on edges per node are set such to keep only the highest intensity edges in the visualization should the settings imply edge overflow. Such settings make sense in the context of the tool; when trying to propagate annotations one does not require hundreds or thousands of edges to some node, but rather the best matches in the local topology.

Alternative approaches that could have been employed to reduce dense “hairball” network drawings commonly use some form of summarization [64]. One possible solution is to (hierarchically) aggregate nodes based on attribute or topological similarity [65], i.e. group sets of related nodes into so-called hypernodes in order to only render these hypernodes and edges between them. Alternatively, edge connections could be simplified, using so-called edge-bundling, to reduce visual clutter and allow for a better exploration of the network’s global topology [66]. However, by introducing such level-of-detail hierarchies, summarization manipulates the network’s topology [67], thereby altering the perceived relationships within the network and potentially confusing the mental map created by the user. Importantly, no matter how information is aggregated, such representations must be combined with meaningful, and potentially complex interaction techniques, such as hypernode expansion or semantic zooming [68, 69], to allow users to effectively explore networks at different levels of granularity. No matter the approach taken, some form of data reduction or simplification via filters needs to be used with interactive elements unlocking additional details on demand. SpecXplore, however, aims to only minimally alter the perceived topology of the network and provide intuitive and easily understandable forms of interaction.

### 4.3 k-medoid clustering in specXplore

SpecXplore makes use of k-medoid clustering as a means of providing feature groupings that complement the t-SNE embedding with a more concrete sense of local clustering via group-based color highlighting. k-medoid has definite computational and interpretational advantages over clustering approaches such as K-means as it avoids the need to compute centroids and recompute distances to the latter. Here, rather than computing centroids for each cluster, and recomputing distances between those centroids and the cluster member features, k-medoid clustering makes use of features with median distance from all other features in the cluster as medoids against which to measure average distances. This makes k-medoid clustering suitable for use with any pairwise distance matrix and allows it to avoid potentially costly distance computations. It also means that t-SNE embeddings based on ms2deepscore pairwise similarities can be complemented by the additional highlighting of local clusters independent of t-SNE 2D projection artefacts.

We considered making use of agglomerative or divisive hierarchical clustering approaches making use of medoids in linkage to maintain consistency between different levels of granularity in clustering [70]. This would have a mental map advantage as clusters would have constrained relationships between levels of granularity, i.e. a larger cluster would contain a number of smaller clusters, each themselves containing a number of smaller clusters. For k-medoid clustering, no such agreement between different values of K is enforced, and hence cluster assignments may vary across settings of K sometimes grouping features together and sometimes not. However, we ultimately opted against making use of hierarchical clustering in specXplore for two reasons. First, implementations of hierarchical approaches using medoids are few, and not available in our Python environment. Second, at any level of hierarchy, clustering would be sub-optimal when compared to single level clustering because of superimposed hierarchy constraints [70]. Thinking of similarity dependent clustering as an assistive grouping in the t-SNE embedding to be superseded with partial network drawing based explorations using thresholds, we opted to keep clustering as simple as possible, merely giving possible starting subdivisions of the data for various k.

More work on measuring and comparing optimality would be needed to provide users with stronger guidelines for automatic algorithmic clustering. An interesting additional research angle requiring formal and automated clustering guidelines we are currently considering is to make use of kmedoid clustering as a means of storing, propagating, and re-calibrating chemical annotations in a group-wise and automated setting. Such automated efforts could naturally also be channeled back into specXplore, providing users with additional information for use in their annotation efforts.

## 5 Conclusion

SpecXplore provides a means of interactively slicing into the complete matrix of pairwise spectral similarities via adjustable settings and complementary views. Exploration of the data in this way provides a means of determining spectral neighborhoods of interest and to assist direct and indirect network annotation propagation. In addition, specXplore exposes the impact of topological filtering settings on effective topology and local neighborhood contexts. Being based on ms2deepscore, it further allows for state of the art similarity scoring that reflects structural similarities more closely. This in turn opens up opportunities for finding similarities missed by traditional scoring approaches. SpecXplore thus provides users with flexible state of the art data exploration platform. Future work for specXplore we are considering are a) providing an online hosted version for more straightforward accessibility, and b) an integration with statistical testing results for more effective prioritization levering experimental designs (such as in FERMO [71]).

## 6 Acknowledgements

The wheat dataset was kindly provided Rainer Schumacher and the BOKU Core Facility Bioactive Molecules: Screening and Analysis.

## Bibliography

[1] Niek F. de Jonge, Kevin Mildau, David Meijer, Joris J. R. Louwen, Christoph Bueschl, Florian Huber, and Justin J. J. van der Hooft. Good practices and recommendations for using and benchmarking computational metabolomics metabolite annotation tools. Metabolomics, 18(12):103, Dec 2022. ISSN 1573-3890. doi: 10.1007/s11306-022-01963-y. URL 10.1007/s11306-022-01963-y.

[2] Adam Amara, Clément Frainay, Fabien Jourdan, Thomas Naake, Steffen Neumann, Elva María Novoa-del Toro, Reza M Salek, Liesa Salzer, Sarah Scharfenberg, and Michael Witting. Networks and graphs discovery in metabolomics data analysis and interpretation. Frontiers in Molecular Biosciences, 9, 2022. ISSN 2296-889X. doi: 10.3389/fmolb.2022.841373. URL https://www.frontiersin.org/articles/10.3389/fmolb.2022.841373.

[3] Jeramie Watrous, Patrick Roach, Theodore Alexandrov, Brandi S. Heath, Jane Y. Yang, Roland D. Kersten, Menno van der Voort, Kit Pogliano, Harald Gross, Jos M. Raaijmakers, Bradley S. Moore, Julia Laskin, Nuno Bandeira, and Pieter C. Dorrestein. Mass spectral molecular networking of living microbial colonies. Proceedings of the National Academy of Sciences, 109(26):E1743–E1752, 2012. doi: 10.1073/pnas.1203689109. URL 10.1073/pnas.1203689109.

[4] Mingxun Wang, Jeremy J. Carver, Vanessa V. Phelan, Laura M. Sanchez, Neha Garg, Yao Peng, Don Duy Nguyen, Jeramie Watrous, Clifford A. Kapono, Tal Luzzatto-Knaan, Carla Porto, Amina Bouslimani, Alexey V. Melnik, Michael J. Meehan, Wei-Ting Liu, Max Crüsemann, Paul D. Boudreau, Eduardo Esquenazi, Mario Sandoval-Calderón, Roland D. Kersten, Laura A. Pace, Robert A. Quinn, Katherine R. Duncan, Cheng-Chih Hsu, Dimitrios J. Floros, Ronnie G. Gavilan, Karin Kleigrewe, Trent Northen, Rachel J. Dutton, Delphine Parrot, Erin E. Carlson, Bertrand Aigle, Charlotte F. Michelsen, Lars Jelsbak, Christian Sohlenkamp, Pavel Pevzner, Anna Edlund, Jeffrey McLean, Jörn Piel, Brian T. Murphy, Lena Gerwick, Chih-Chuang Liaw, Yu-Liang Yang, Hans-Ulrich Humpf, Maria Maansson, Robert A. Keyzers, Amy C. Sims, Andrew R. Johnson, Ashley M. Side-bottom, Brian E. Sedio, Andreas Klitgaard, Charles B. Larson, Cristopher A. Boya P, Daniel Torres-Mendoza, David J. Gonzalez, Denise B. Silva, Lucas M. Marques, Daniel P. Demarque, Egle Pociute, Ellis C. O’Neill, Enora Briand, Eric J. N. Helfrich, Eve A. Granatosky, Evgenia Glukhov, Florian Ryffel, Hailey Houson, Hosein Mohimani, Jenan J. Kharbush, Yi Zeng, Julia A. Vorholt, Kenji L. Kurita, Pep Charusanti, Kerry L. McPhail, Kristian Fog Nielsen, Lisa Vuong, Maryam Elfeki, Matthew F. Traxler, Niclas Engene, Nobuhiro Koyama, Oliver B. Vining, Ralph Baric, Ricardo R. Silva, Samantha J. Mascuch, Sophie Tomasi, Stefan Jenkins, Venkat Macherla, Thomas Hoffman, Vinayak Agarwal, Philip G. Williams, Jingqui Dai, Ram Neupane, Joshua Gurr, Andrés M. C. Rodríguez, Anne Lamsa, Chen Zhang, Kathleen Dorrestein, Brendan M. Duggan, Jehad Almaliti, Pierre-Marie Allard, Prasad Phapale, Louis-Felix Nothias, Theodore Alexandrov, Marc Litaudon, Jean-Luc Wolfender, Jennifer E. Kyle, Thomas O. Metz, Tyler Peryea, Dac-Trung Nguyen, Danielle VanLeer, Paul Shinn, Ajit Jadhav, Rolf Müller, Katrina M. Waters, Wenyuan Shi, Xueting Liu, Lixin Zhang, Rob Knight, Paul R. Jensen, Bernhard Ø Palsson, Kit Pogliano, Roger G. Linington, Marcelino Gutiérrez, Norberto P. Lopes, William H. Gerwick, Bradley S. Moore, Pieter C. Dorrestein, and Nuno Bandeira. Sharing and community curation of mass spectrometry data with global natural products social molecular networking. Nature Biotechnology, 34(8):828–837, Aug 2016. ISSN 1546-1696. doi: 10.1038/nbt.3597. URL 10.1038/nbt.3597.

[5] Louis-Félix Nothias, Daniel Petras, Robin Schmid, Kai Dührkop, Johannes Rainer, Abinesh Sarvepalli, Ivan Protsyuk, Madeleine Ernst, Hiroshi Tsugawa, Markus Fleischauer, Fabian Aicheler, Alexander A. Aksenov, Oliver Alka, Pierre-Marie Allard, Aiko Barsch, Xavier Cachet, Andres Mauricio Caraballo-Rodriguez, Ricardo R. Da Silva, Tam Dang, Neha Garg, Julia M. Gauglitz, Alexey Gurevich, Giorgis Isaac, Alan K. Jarmusch, Zdeněk Kameník, Kyo Bin Kang, Nikolas Kessler, Irina Koester, Ansgar Korf, Audrey Le Gouellec, Marcus Ludwig, Christian Martin H., Laura-Isobel McCall, Jonathan McSayles, Sven W. Meyer, Hosein Mohimani, Mustafa Morsy, Oriane Moyne, Steffen Neumann, Heiko Neuweger, Ngoc Hung Nguyen, Melissa Nothias-Esposito, Julien Paolini, Vanessa V. Phelan, Tomáš Pluskal, Robert A. Quinn, Simon Rogers, Bindesh Shrestha, Anupriya Tripathi, Justin J. J. van der Hooft, Fernando Vargas, Kelly C. Weldon, Michael Witting, Heejung Yang, Zheng Zhang, Florian Zubeil, Oliver Kohlbacher, Sebastian Böcker, Theodore Alexandrov, Nuno Bandeira, Mingxun Wang, and Pieter C. Dorrestein. Feature-based molecular networking in the gnps analysis environment. Nature Methods, 17(9):905–908, Sep 2020. ISSN 1548-7105. doi: 10.1038/s41592-020-0933-6. URL 10.1038/s41592-020-0933-6.

[6] Louis-Félix Nothias, Mélissa Nothias-Esposito, Ricardo da Silva, Mingxun Wang, Ivan Protsyuk, Zheng Zhang, Abi Sarvepalli, Pieter Leyssen, David Touboul, Jean Costa, Julien Paolini, Theodore Alexandrov, Marc Litaudon, and Pieter C. Dorrestein. Bioactivity-based molecular networking for the discovery of drug leads in natural product bioassay-guided fractionation. Journal of Natural Products, 81(4):758–767, 2018. doi: 10.1021/acs.jnatprod.7b00737. URL 10.1021/acs.jnatprod.7b00737. PMID: 29498278.

[7] Alexander E. Fox Ramos, Laurent Evanno, Erwan Poupon, Pierre Champy, and Mehdi A. Beniddir. Natural products targeting strategies involving molecular networking: different manners, one goal. Nat. Prod. Rep., 36: 960–980, 2019. doi: 10.1039/C9NP00006B. URL 10.1039/C9NP00006B.

[8] Guo-Fei Qin, Xiao Zhang, Feng Zhu, Zong-Qing Huo, Qing-Qiang Yao, Qun Feng, Zhong Liu, Gui-Min Zhang, Jing-Chun Yao, and Hong-Bao Liang. Ms/ms-based molecular networking: An efficient approach for natural products dereplication. Molecules, 28(1), 2023. ISSN 1420-3049. doi: 10.3390/molecules28010157. URL https://www.mdpi.com/1420-3049/28/1/157.

[9] Zhitao Tian, Fangzhou Liu, Dongqin Li, Alisdair R. Fernie, and Wei Chen. Strategies for structure elucidation of small molecules based on lc–ms/ms data from complex biological samples. Computational and Structural Biotechnology Journal, 20:5085–5097, 2022. ISSN 2001-0370. doi: 10.1016/j.csbj.2022.09.004. URL https://www.sciencedirect.com/science/article/pii/S2001037022004093.

[10] Florian Huber, Sven van der Burg, Justin J. J. van der Hooft, and Lars Ridder. Ms2deepscore: a novel deep learning similarity measure to compare tandem mass spectra. Journal of Cheminformatics, 13(1):84, Oct 2021. ISSN 1758-2946. doi: 10.1186/s13321-021-00558-4. URL 10.1186/s13321-021-00558-4.

[11] Robin Schmid, Steffen Heuckeroth, Ansgar Korf, Aleksandr Smirnov, Owen Myers, Thomas S. Dyrlund, Roman Bushuiev, Kevin J. Murray, Nils Hoffmann, Miaoshan Lu, Abinesh Sarvepalli, Zheng Zhang, Markus Fleischauer, Kai Dührkop, Mark Wesner, Shawn J. Hoogstra, Edward Rudt, Olena Mokshyna, Corinna Brungs, Kirill Ponomarov, Lana Mutabdžija, Tito Damiani, Chris J. Pudney, Mark Earll, Patrick O. Helmer, Timothy R. Fallon, Tobias Schulze, Albert Rivas-Ubach, Aivett Bilbao, Henning Richter, Louis-Félix Nothias, Mingxun Wang, Matej Orešič, Jing-Ke Weng, Sebastian Böcker, Astrid Jeibmann, Heiko Hayen, Uwe Karst, Pieter C. Dorrestein, Daniel Petras, Xiuxia Du, and Tomáš Pluskal. Integrative analysis of multimodal mass spectrometry data in MZmine 3. Nature Biotechnology, 41(4):447–449, March 2023. doi: 10.1038/s41587-023-01690-2. URL 10.1038/s41587-023-01690-2.

[12] Florian Huber, Stefan Verhoeven, Christiaan Meijer, Hanno Spreeuw, Efraín Manuel Villanueva Castilla, Cunliang Geng, Justin J. j. van der Hooft, Simon Rogers, Adam Belloum, Faruk Diblen, and Jurriaan H. Spaaks. matchms - processing and similarity evaluation of mass spectrometry data. Journal of Open Source Software, 5(52):2411, 2020. doi: 10.21105/joss.02411. URL 10.21105/joss.02411.

[13] Florian Huber, Lars Ridder, Stefan Verhoeven, Jurriaan H. Spaaks, Faruk Diblen, Simon Rogers, and Justin J. J. van der Hooft. Spec2vec: Improved mass spectral similarity scoring through learning of structural relationships. PLOS Computational Biology, 17(2):1–18, 02 2021. doi: 10.1371/journal.pcbi.1008724. URL 10.1371/journal.pcbi.1008724.

[14] Niek F. de Jonge, Joris R. Louwen, Elena Chekmeneva, Stephane Camuzeaux, Femke J. Vermeir, Robert S. Jansen, Florian Huber, and Justin J.J. van der Hooft. Ms2query: Reliable and scalable ms2 mass spectral-based analogue search. bioRxiv, 2022. doi: 10.1101/2022.07.22.501125. URL https://www.biorxiv.org/content/early/2022/07/23/2022.07.22.501125.

[15] Laurens van der Maaten and Geoffrey Hinton. Visualizing data using t-sne. Journal of Machine Learning Research, 9(86):2579–2605, 2008. URL http://jmlr.org/papers/v9/vandermaaten08a.html.

[16] F. Pedregosa, G. Varoquaux, A. Gramfort, V. Michel, B. Thirion, O. Grisel, M. Blondel, P. Prettenhofer, R. Weiss, V. Dubourg, J. Vanderplas, A. Passos, D. Cournapeau, M. Brucher, M. Perrot, and E. Duchesnay. Scikit-learn: Machine learning in Python. Journal of Machine Learning Research, 12:2825–2830, 2011.

[17] Angelos Chatzimparmpas, Rafael Messias Martins, and Andreas Kerren. t-visne: A visual inspector for the exploration of t-sne. In IEEE Information Visualization (VIS’18), Berlin, Germany, 21-26 October, 2018, 2018.

[18] Erich Schubert and Peter J. Rousseeuw. Faster k-medoids clustering: Improving the pam, clara, and clarans algorithms. In Giuseppe Amato, Claudio Gennaro, Vincent Oria, and Miloš Radovanović, editors, Similarity Search and Applications, pages 171–187, Cham, 2019. Springer International Publishing. ISBN 978-3-030-32047-8.

[19] Kai Dührkop, Markus Fleischauer, Marcus Ludwig, Alexander A. Aksenov, Alexey V. Melnik, Marvin Meusel, Pieter C. Dorrestein, Juho Rousu, and Sebastian Böcker. Sirius 4: a rapid tool for turning tandem mass spectra into metabolite structure information. Nature Methods, 16(4):299–302, Apr 2019. ISSN 1548-7105. doi: 10.1038/s41592-019-0344-8. URL 10.1038/s41592-019-0344-8.

[20] Kai Dührkop, Louis-Félix Nothias, Markus Fleischauer, Raphael Reher, Marcus Ludwig, Martin A Hoffmann, Daniel Petras, William H Gerwick, Juho Rousu, Pieter C Dorrestein, and Sebastian Böcker. Systematic classification of unknown metabolites using high-resolution fragmentation mass spectra. Nat Biotechnol, 39(4):462–471, November 2020.

[21] Ji Soo Yi, Youn ah Kang, John Stasko, and J.A. Jacko. Toward a deeper understanding of the role of interaction in information visualization. IEEE Transactions on Visualization and Computer Graphics, 13(6):1224–1231, 2007. doi: 10.1109/TVCG.2007.70515.

[22] Shammamah Hossain. Visualization of Bioinformatics Data with Dash Bio. In Chris Calloway, David Lippa, Dillon Niederhut, and David Shupe, editors, Proceedings of the 18th Python in Science Conference, pages 126 –133, 2019. doi: 10.25080/Majora-7ddc1dd1-012.

[23] Max Franz, Christian T. Lopes, Gerardo Huck, Yue Dong, Onur Sumer, and Gary D. Bader. Cytoscape.js: a graph theory library for visualisation and analysis. Bioinformatics, 32(2):309–311, sep 2015. doi: 10.1093/bioinformatics/btv557. URL 10.1093%2Fbioinformatics%2Fbtv557.

[24] Max Franz, Christian T Lopes, Dylan Fong, Mike Kucera, Manfred Cheung, Metin Can Siper, Gerardo Huck, Yue Dong, Onur Sumer, and Gary D Bader. Cytoscape.js 2023 update: a graph theory library for visualization and analysis. Bioinformatics, 39(1), jan 2023. doi: 10.1093/bioinformatics/btad031. URL 10.1093%2Fbioinformatics%2Fbtad031.

[25] Tamara Munzner. Visualization Analysis and Design. A K Peters/CRC Press, New York, October 2014. ISBN 978-0-429-08890-2. doi: 10.1201/b17511.

[26] Christopher G Healey. Choosing effective colours for data visualization. In Proceedings of Seventh Annual IEEE Visualization’96, pages 263–270. IEEE, 1996.

[27] Ben Shneiderman. The Eyes Have It: A Task by Data Type Taxonomy for Information Visualizations. In BENJAMIN B. Bederson and BEN Shneiderman, editors, The Craft of Information Visualization, Interactive Technologies, pages 364–371. Morgan Kaufmann, San Francisco, January 2003. ISBN 978-1-55860-915-0. doi: 10.1016/B978-155860915-0/50046-9. URL https://www.sciencedirect.com/science/article/pii/B9781558609150500469.

[28] Wesley Willett, Jeffrey Heer, and Maneesh Agrawala. Scented Widgets: Improving Navigation Cues with Embedded Visualizations. IEEE Transactions on Visualization and Computer Graphics, 13(6):1129–1136, November 2007. ISSN 1941-0506. doi: 10.1109/TVCG.2007.70589. Conference Name: IEEE Transactions on Visualization and Computer Graphics.

[29] Yanhong Wu, Naveen Pitipornvivat, Jian Zhao, Sixiao Yang, Guowei Huang, and Huamin Qu. egoSlider: Visual Analysis of Egocentric Network Evolution. IEEE Transactions on Visualization and Computer Graphics, 22(1):260–269, January 2016. ISSN 1941-0506. doi: 10.1109/TVCG.2015.2468151. Conference Name: IEEE Transactions on Visualization and Computer Graphics.

[30] Lei Shi, Chen Wang, and Zhen Wen. Dynamic network visualization in 1.5D. In 2011 IEEE Pacific Visualization Symposium, pages 179–186, March 2011. doi: 10.1109/PACIFICVIS.2011.5742388. ISSN: 2165-8773.

[31] Maria Doppler, Christoph Bueschl, Bernhard Kluger, Andrea Koutnik, Marc Lemmens, Hermann Buerstmayr, Justyna Rechthaler, Rudolf Krska, Gerhard Adam, and Rainer Schuhmacher. Stable isotope–assisted plant metabolomics: Combination of global and tracer-based labeling for enhanced untargeted profiling and compound annotation. Frontiers in Plant Science, 10:1366, 2019.

[32] Christoph Bueschl, Maria Doppler, Elisabeth Varga, Bernhard Seidl, Mira Flasch, Benedikt Warth, and Juergen Zanghellini. Peakbot: machine-learning-based chromatographic peak picking. Bioinformatics, 38(13):3422–3428, 2022.

[33] Ian Oesterle, Manuel Pristner, Sabrina Berger, Mingxun Wang, Vinicius Verri Hernandes, Annette Rompel, and Benedikt Warth. Exposomic biomonitoring of polyphenols by non-targeted analysis and suspect screening. Analytical Chemistry, 95(28):10686–10694, 2023.

[34] Madeleine Ernst, Kyo Bin Kang, Andrés Mauricio Caraballo-Rodríguez, Louis-Felix Nothias, Joe Wandy, Christopher Chen, Mingxun Wang, Simon Rogers, Marnix H. Medema, Pieter C. Dorrestein, and Justin J.J. van der Hooft. Molnetenhancer: Enhanced molecular networks by integrating metabolome mining and annotation tools. Metabolites, 9(7), 2019. ISSN 2218-1989. doi: 10.3390/metabo9070144. URL https://www.mdpi.com/2218-1989/9/7/144.

[35] Florent Olivon, Nicolas Elie, Gwendal Grelier, Fanny Roussi, Marc Litaudon, and David Touboul. Metgem software for the generation of molecular networks based on the t-sne algorithm. Analytical Chemistry, 90(23): 13900–13908, 2018. doi: 10.1021/acs.analchem.8b03099. URL 10.1021/acs.analchem.8b03099. PMID: 30335965.

[36] Nicolas Elie, Cyrille Santerre, and David Touboul. Generation of a molecular network from electron ionization mass spectrometry data by combining mzmine2 and metgem software. Analytical Chemistry, 91(18):11489–11492, 2019. doi: 10.1021/acs.analchem.9b02802. URL 10.1021/acs.analchem.9b02802. PMID: 31429549.

[37] Justin Johan Jozias van der Hooft, Joe Wandy, Michael P. Barrett, Karl E. V. Burgess, and Simon Rogers. Topic modeling for untargeted substructure exploration in metabolomics. Proceedings of the National Academy of Sciences, 113(48):13738–13743, 2016. doi: 10.1073/pnas.1608041113. URL https://www.pnas.org/doi/abs/10.1073/pnas.1608041113.

[38] Paul Shannon, Andrew Markiel, Owen Ozier, Nitin S Baliga, Jonathan T Wang, Daniel Ramage, Nada Amin, Benno Schwikowski, and Trey Ideker. Cytoscape: a software environment for integrated models of biomolecular interaction networks. Genome Res, 13(11):2498–2504, November 2003.

[39] Yongyi Li, Zhirong Cui, Ying Li, Juanjuan Gao, Rong Tao, Jixin Li, Yi Li, and Jun Luo. Integrated molecular networking strategy enhance the accuracy and visualization of components identification: A case study of ginkgo biloba leaf extract. Journal of Pharmaceutical and Biomedical Analysis, 209:114523, 2022. ISSN 0731-7085. doi: j.jpba.2021.114523. URL https://www.sciencedirect.com/science/article/pii/S0731708521006348.

[40] Lerato Nephali, Paul Steenkamp, Karl Burgess, Johan Huyser, Margaretha Brand, Justin J. J. van der Hooft, and Fidele Tugizimana. Mass spectral molecular networking to profile the metabolome of biostimulant bacillus strains. Frontiers in Plant Science, 13, 2022. ISSN 1664-462X. doi: 10.3389/fpls.2022.920963. URL https://www.frontiersin.org/articles/10.3389/fpls.2022.920963.

[41] Morena M. Tinte, Keabetswe Masike, Paul A. Steenkamp, Johan Huyser, Justin J. J. van der Hooft, and Fidele Tugizimana. Computational metabolomics tools reveal metabolic reconfigurations underlying the effects of biostimulant seaweed extracts on maize plants under drought stress conditions. Metabolites, 12(6), 2022. ISSN 2218-1989. doi: 10.3390/metabo12060487. URL https://www.mdpi.com/2218-1989/12/6/487.

[42] Téo Hebra, Nicolas Elie, Salomé Poyer, Elsa Van Elslande, David Touboul, and Véronique Eparvier. Dereplication, annotation, and characterization of 74 potential antimicrobial metabolites from penicillium sclerotiorum using t-sne molecular networks. Metabolites, 11(7), 2021. ISSN 2218-1989. doi: 10.3390/metabo11070444. URL https://www.mdpi.com/2218-1989/11/7/444.

[43] Téo Hebra, Nicolas Pollet, David Touboul, and Véronique Eparvier. Combining osmac, metabolomic and genomic methods for the production and annotation of halogenated azaphilones and ilicicolins in termite symbiotic fungi. Scientific Reports, 12(1):17310, Oct 2022. ISSN 2045-2322. doi: 10.1038/s41598-022-22256-3. URL 10.1038/s41598-022-22256-3.

[44] Jonathan Sorres, Téo Hebra, Nicolas Elie, Charlotte Leman-Loubière, Tatyana Grayfer, Philippe Grellier, David Touboul, Didier Stien, and Véronique Eparvier. Antiparasitic ovalicin derivatives from pseudallescheria boydii, a mutualistic fungus of french guiana termites. Molecules, 27(4), 2022. ISSN 1420-3049. doi: 10.3390/molecules27041182. URL https://www.mdpi.com/1420-3049/27/4/1182.

[45] Marian Dörk, Sheelagh Carpendale, and Carey Williamson. EdgeMaps: visualizing explicit and implicit relations. In Visualization and Data Analysis 2011. SPIE, January 2011. doi: 10.1117/12.872578. URL 10.1117/12.872578.

[46] M. Dork, S. Carpendale, and C. Williamson. Visualizing explicit and implicit relations of complex information spaces. Information Visualization, 11(1):5–21, November 2011. doi: 10.1177/1473871611425872. URL 10.1177/1473871611425872.

[47] Ricardo R. da Silva, Mingxun Wang, Louis-Félix Nothias, Justin J. J. van der Hooft, Andrés Mauricio Caraballo-Rodríguez, Evan Fox, Marcy J. Balunas, Jonathan L. Klassen, Norberto Peporine Lopes, and Pieter C. Dorrestein. Propagating annotations of molecular networks using in silico fragmentation. PLOS Computational Biology, 14 (4):1–26, 04 2018. doi: 10.1371/journal.pcbi.1006089. URL 10.1371/journal.pcbi.1006089.

[48] Nicholas J. Morehouse, Trevor N. Clark, Emily J. McMann, Jeffrey A. van Santen, F. P. Jake Haeckl, Christopher A. Gray, and Roger G. Linington. Annotation of natural product compound families using molecular networking topology and structural similarity fingerprinting. Nature Communications, 14(1):308, Jan 2023. ISSN 2041-1723. doi: 10.1038/s41467-022-35734-z. URL 10.1038/s41467-022-35734-z.

[49] Bongshin Lee, Catherine Plaisant, Cynthia Sims Parr, Jean-Daniel Fekete, and Nathalie Henry. Task taxonomy for graph visualization. In Proceedings of the 2006 AVI Workshop on BEyond Time and Errors: Novel Evaluation Methods for Information Visualization, BELIV ‘06, page 1–5, New York, NY, USA, 2006. Association for Computing Machinery. ISBN 1595935622. doi: 10.1145/1168149.1168168. URL 10.1145/1168149.1168168.

[50] Wout Bittremieux, Robin Schmid, Florian Huber, Justin J. J. van der Hooft, Mingxun Wang, and Pieter C. Dorrestein. Comparison of cosine, modified cosine, and neutral loss based spectrum alignment for discovery of structurally related molecules. Journal of the American Society for Mass Spectrometry, 33(9):1733–1744, 2022. doi: 10.1021/jasms.2c00153. URL 10.1021/jasms.2c00153. PMID: 35960544.

[51] Yuanyue Li, Tobias Kind, Jacob Folz, Arpana Vaniya, Sajjan Singh Mehta, and Oliver Fiehn. Spectral entropy outperforms ms/ms dot product similarity for small-molecule compound identification. Nature Methods, 18(12): 1524–1531, Dec 2021. ISSN 1548-7105. doi: 10.1038/s41592-021-01331-z. URL 10.1038/s41592-021-01331-z.

[52] Eades P. A heuristic for graph drawing. Congressus Numerantium, 42:149–160, 1984. URL https://cir.nii.ac.jp/crid/1573387448853684864.

[53] Thomas M. J. Fruchterman and Edward M. Reingold. Graph drawing by force-directed placement. Software: Practice and Experience, 21(11):1129–1164, 1991. ISSN 1097-024X. doi: 10.1002/spe.4380211102. URL https://onlinelibrary.wiley.com/doi/abs/10.1002/spe.4380211102. eprint: https://onlinelibrary.wiley.com/doi/pdf/10.1002/spe.4380211102.

[54] Tomihisa Kamada and Satoru Kawai. AN ALGORITHM FOR DRAWING GENERAL UNDIRECTED GRAPHS. INFORMATION PROCESSING LETTERS, 31(1):9, 1989.

[55] Stephen G. Kobourov. Spring Embedders and Force Directed Graph Drawing Algorithms. arXiv:1201.3011 [cs], January 2012. URL http://arxiv.org/abs/1201.3011. xarXiv: 1201.3011.

[56] Helen Purchase. Which aesthetic has the greatest effect on human understanding? In Giuseppe DiBattista, editor, Graph Drawing, Lecture Notes in Computer Science, pages 248–261, Berlin, Heidelberg, 1997. Springer. ISBN 978-3-540-69674-2. doi: 10.1007/3-540-63938-167.

[57] Helen C. Purchase, David Carrington, and Jo-Anne Allder. Empirical Evaluation of Aesthetics-based Graph Layout. Empirical Software Engineering, 7(3):233–255, September 2002. ISSN 1573-7616. doi: 10.1023/A:1016344215610. URL 10.1023/A:1016344215610.

[58] Helen C. Purchase, Christopher Pilcher, and Beryl Plimmer. Graph Drawing Aesthetics—Created by Users, Not Algorithms. IEEE Transactions on Visualization and Computer Graphics, 18(1):81–92, January 2012. ISSN 1941-0506. doi: 10.1109/TVCG.2010.269. Conference Name: IEEE Transactions on Visualization and Computer Graphics.

[59] Stephen G. Kobourov, Sergey Pupyrev, and Bahador Saket. Are Crossings Important for Drawing Large Graphs? In Eduardo Bayro-Corrochano and Edwin Hancock, editors, Progress in Pattern Recognition, Image Analysis, Computer Vision, and Applications, volume 8827, pages 234–245. Springer International Publishing, Cham, 2014. ISBN 978-3-319-12567-1 978-3-319-12568-8. doi: 10.1007/978-3-662-45803-720. URL http://link.springer.com/10.1007/978-3-662-45803-7_20. Series Title: Lecture Notes in Computer Science.

[60] James Abello, Daniel Mawhirter, and Kevin Sun. Taming a Graph Hairball: Local Exploration in a Global Context. In Pablo Moscato and Natalie Jane de Vries, editors, Business and Consumer Analytics: New Ideas, pages 467–490. Springer International Publishing, Cham, 2019. ISBN 978-3-030-06222-4. doi: 10.1007/978-3-030-06222-410. URL 10.1007/978-3-030-06222-4_10.

[61] Arlind Nocaj, Mark Ortmann, and Ulrik Brandes. Untangling Hairballs. In Christian Duncan and Antonios Symvonis, editors, Graph Drawing, Lecture Notes in Computer Science, pages 101–112, Berlin, Heidelberg, 2014. Springer. ISBN 978-3-662-45803-7. doi: 10.1007/978-3-662-45803-79.

[62] Juuso Parkkinen, Kristian Nybo, Jaakko Peltonen, and Samuel Kaski. Graph visualization with latent variable models. In Proceedings of the Eighth Workshop on Mining and Learning with Graphs, MLG ‘10, pages 94–101, New York, NY, USA, July 2010. Association for Computing Machinery. ISBN 978-1-4503-0214-2. doi: 10.1145/1830252.1830265. URL 10.1145/1830252.1830265.

[63] Till Bruckdorfer, Sabine Cornelsen, Carsten Gutwenger, Michael Kaufmann, Fabrizio Montecchiani, Martin Nöllenburg, and Alexander Wolff. Progress on Partial Edge Drawings. In Walter Didimo and Maurizio Patrignani, editors, Graph Drawing, Lecture Notes in Computer Science, pages 67–78, Berlin, Heidelberg, 2013. Springer. ISBN 978-3-642-36763-2. doi: 10.1007/978-3-642-36763-27.

[64] Yike Liu, Tara Safavi, Abhilash Dighe, and Danai Koutra. Graph Summarization Methods and Applications: A Survey. ACM Computing Surveys, 51(3):62:1–62:34, June 2018. ISSN 0360-0300. doi: 10.1145/3186727. URL 10.1145/3186727.

[65] Niklas Elmqvist and Jean-Daniel Fekete. Hierarchical Aggregation for Information Visualization: Overview, Techniques, and Design Guidelines. IEEE Transactions on Visualization and Computer Graphics, 16(3):439–454, May 2010. ISSN 1941-0506. doi: 10.1109/TVCG.2009.84. Conference Name: IEEE Transactions on Visualization and Computer Graphics.

[66] Hong Zhou, Panpan Xu, Xiaoru Yuan, and Huamin Qu. Edge bundling in information visualization. Tsinghua Science and Technology, 18(2):145–156, April 2013. ISSN 1007-0214. doi: 10.1109/TST.2013.6509098. Conference Name: Tsinghua Science and Technology.

[67] Kathryn Gray, Mingwei Li, Reyan Ahmed, Md Khaledur Rahman, Ariful Azad, Stephen Kobourov, and Katy Börner. A scalable method for readable tree layouts. IEEE Transactions on Visualization and Computer Graphics, 2023.

[68] Vitalis Wiens, Steffen Lohmann, and Sören Auer. Semantic Zooming for Ontology Graph Visualizations. In Proceedings of the Knowledge Capture Conference, K-CAP 2017, pages 1–8, New York, NY, USA, December 2017. Association for Computing Machinery. ISBN 978-1-4503-5553-7. doi: 10.1145/3148011.3148015. URL 10.1145/3148011.3148015.

[69] Ana Figueiras. Towards the Understanding of Interaction in Information Visualization. In 2015 19th International Conference on Information Visualisation, pages 140–147, July 2015. doi: 10.1109/iV.2015.34. ISSN: 2375-0138.

[70] Erich Schubert. Hacam: Hierarchical agglomerative clustering around medoids-and its limitations. In LWDA, pages 191–204, 2021.

[71] Mitja M. Zdouc, Lina M. Bayona Maldonado, Hannah E. Augustijn, Sylvia Soldatou, Niek de Jonge, Marcel Jaspars, Gilles P. van Wezel, Marnix H. Medema, and Justin J. J. van der Hooft. Fermo: a dashboard for streamlined rationalized prioritization of molecular features from mass spectrometry data. bioRxiv, 2022. doi: 10.1101/2022.12.21.521422. URL https://www.biorxiv.org/content/early/2022/12/22/2022.12.21.521422.

