## Supplementary Information pdf for "Tailored mass spectral data exploration using the specXplore interactive dashboard"

Kevin Mildau [conceptualization, implementation, writing - original draft][1,2,3,*], Henry Ehlers [conceptualization, implementation, writing - original draft][4,*], Ian Oesterle [resources][2,5,6], Manuel Pristner [resources][5], Benedikt Warth [resources][5], Maria Doppler [resources][7], Christoph Bueschl [conceptualization, supervision, writing - review & editing][7], Juergen Zanghellini [funding, conceptualization, supervision, , writing - review & editing][1,**], and Justin J.J van der Hooft [conceptualization, supervision, , writing - review & editing][8,9,**]

[1]Biochemical Network Analysis Lab, Department of Analytical Chemistry, University of Vienna, Vienna, Austria
[2]Doctoral School in Chemistry (DOSCHEM), University of Vienna, Vienna, Austria
[3] Austrian Centre of Industrial Biotechnology (ACIB GmbH), Graz, Austria
[4]Institute of Visual Computing and Human-Centered Technology, TU Wien, Vienna, Austria
[5]Department of Food Chemistry and Toxicology, University of Vienna, Vienna, Austria.
[6]Department of Biophysical Chemistry, University of Vienna, Vienna, Austria
[7]University of Natural Resources and Life Sciences (BOKU), Vienna / Tulln, Austria
[8]Bioinformatics Group, Wageningen University, Wageningen, The Netherlands
[9]Department of Biochemistry, University of Johannesburg, Johannesburg, South Africa
[*]Shared First Authors.
[**]Corresponding Authors / Shared Last Authors

October 3, 2023

### Contents

1 Urine Data Example 2

2 k-medoid clustering with with higher granularity. 4

3 Wheat Data MS/MS Data Processing 4

4 MS/MS data comparison views: fragmap, fragmentation spectra, and mirror plots 5

5 Tabular Metadata Integration 5

6 Spectral data pre-processing 5

7 Usage and tuning of t-SNE as a fixed network layout 6

8 In-silico spike-in standards 6

9 Color as a medium for prioritization and visual emphasis 6

10 Chemical Class Visualization 7

11 Effective visual analysis using scented widgets 7

12 Augmented Matrix Views 7

13 Bibliography 7

### 1 Urine Data Example

In addition to the wheat data example shown in the main manuscript, we have also run specXplore on an urine exposomics dataset [1]. Here, study participants were subjected to low and high polyphenol dietary interventions to study the impact of high polyphenol diets on the urine metabolome. Urine samples were collected in three distinct phases. First, subjects were put on low polyphenol diet for three days, and their urine samples were sampled for 24 hours of the third day. Second, immediately following the three day consecutive low polyphenol diet, a polyphenol rich smoothie was ingested in the morning of day four. Urine was sampled throughout this fourth day while participants continued the otherwise low polyphenol diet. Third, on the day after the high polyphenol smoothie, subjects were consuming a high polyphenol diet of their choosing and urine samples were measured throughout the day. Samples were measured with high-resolution mass spectrometry (HRMS) and processed with MZMine3 [1]. This dataset is substantially larger than the wheat dataset, containing 3818 spectra after specXplore filtering steps. However, as can be seen in Figure 1, local explorations in the larger t-SNE overview still work similarly. Local node groups show fragmentation overlaps and spectral similarity that can be used to find related groups of spectra. Two examples of small selections of features are shown around spike-in reference standards for illustrative purposes. In both cases, fragmentation overview maps show good overlap in characteristic fragment ions cross the selected nodes.

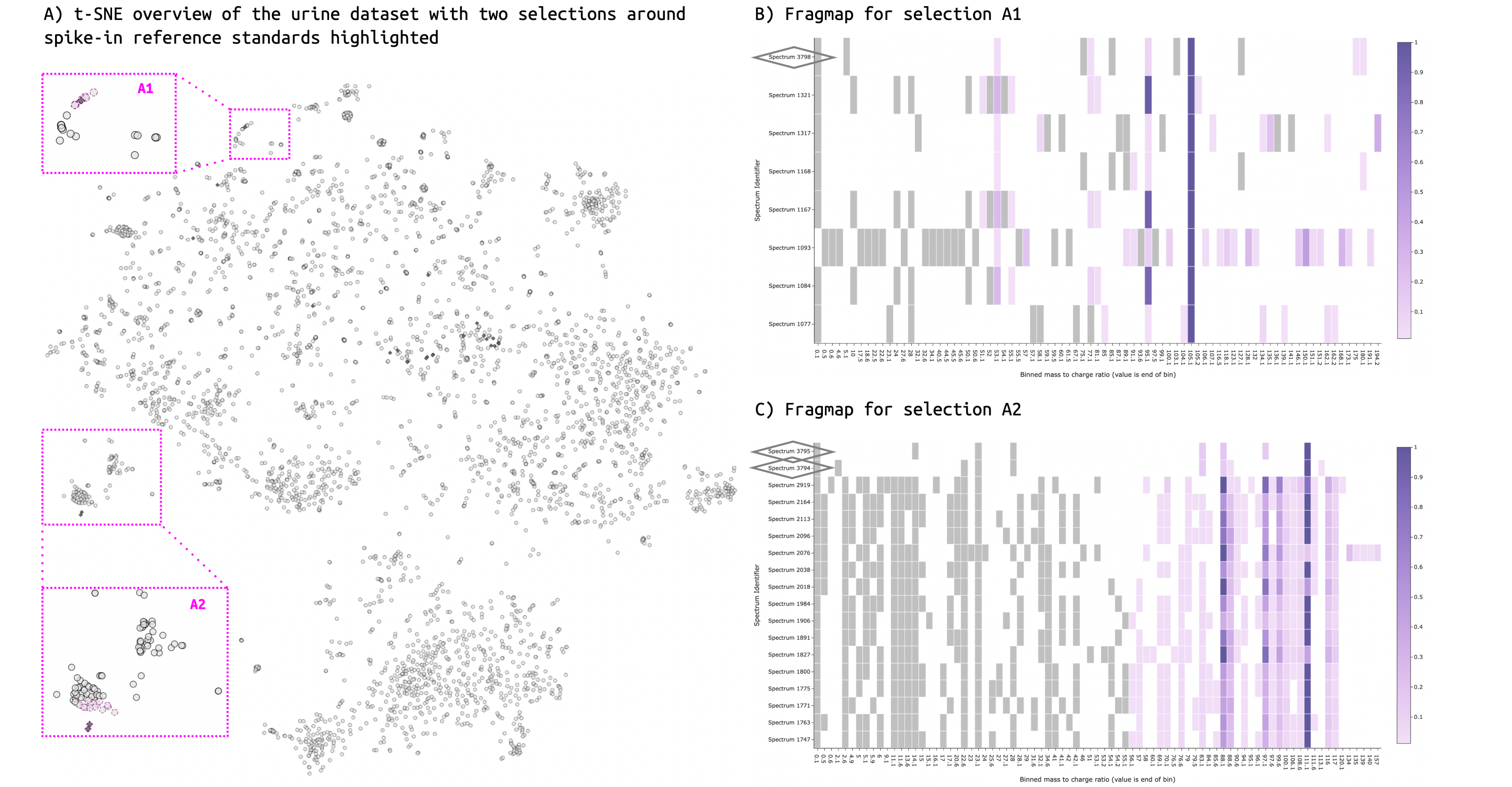


Figure 1: Urine data exploration. A) t-SNE overview representation for the urine dataset. Experimental spectra are represented by circular grey nodes while in-silico spike-in reference standards are highlighted as darker diamond nodes. Two small local node environments involving reference spectra are highlighted. These are in close proximity in the t-SNE embedding and have substantial fragmentation overlap. B) Fragmap for selection A1. There are many fragmentation differences among the spectra. However, high relative intensity fragment ion bins are shared across all spectra in the selection. Spike-in reference spectrum additionally highlighted using diamond outline on fragmap y axis. C) Fragmap for selection A2. Large overlap in binned fragmentation pattern exists between all spectra in this selection. This includes higher intensity fragments, lower intensity fragments, and neutral losses (gray).

#### 2 k-medoid clustering with with higher granularity.

When running k-medoid clustering with larger values of k, the feature groups will tend to become smaller and align well with t-SNE positioning and topological connectivity patterns. Inside specXplore, this can be seen via groupings tending to contain only feature nodes in close proximity to one another in the t-SNE embedding, see wheat data example in Figure 2.

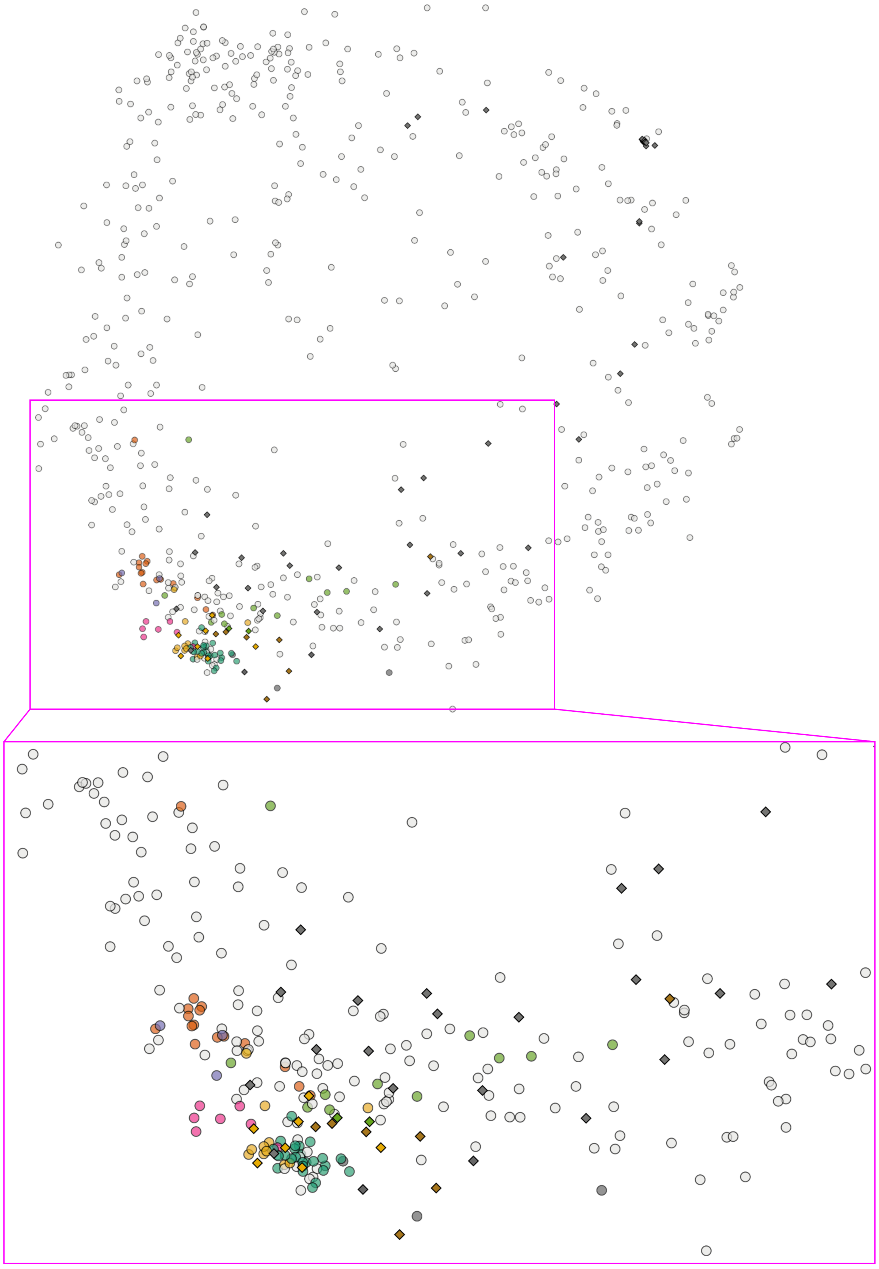

Figure 2: k-medoid clusters in lower left area of the wheat data set example at k = 120. While some node groups are more densely arranged in the t-SNE embedding than others, most groups tend to be highly localized in one area of the embedding.

#### 3 Wheat Data MS/MS Data Processing

All raw files were processed with MZmine 3 (version 3.3.0). Data was loaded in both ionization modes separately [2]. Then mass detection was done with a noise level of 1E4, followed by the ADAP chromatogram builder module (min. number of scans: 4, min. highest intensity: 1E5, scan to scan accuracy: 5 ppm). The chromatograms were smoothed with a Savitzky Golay filter (width 7 points), and chromatographic peaks were detected with the Local minimum feature resolver module (Chromatographic threshold: 10%, minimum absolute height: 1E5, Min. ratio of top to edge: 5, Peak duration: 0.05 – 2 min). The sample specific feature lists were then filtered for only monoisotopic features (Isotopic peaks finder module), and all feature lists were combined using the join aligner module (m/z tolerance: 10 ppm, Retention time tolerance: 0.2 min). Then the combined feature list was filtered for chromatographic peaks with at least one assigned MSMS scans only and all other features were discarded (module Feature list rows filter). Next, the data was exported to the MGF file format using the GNPS export module before further processing with specXplore. Reference standard spectra [1] where spiked into the dataset via inclusion in the MGF file.

### 4 MS/MS data comparison views: fragmap, fragmentation spectra, and mirror plots

SpecXplore makes use of dynamic thresholds in a graph exploration setting. It is hence important to provide the user with insights into whether increased connectivity as a consequence of more liberal thresholds is still reliable or useful. To assist with this, a fragmentation overview heatmap inspired by *UpSet* plots [3] is provided that allows visualizing overlaps in fragmentation between sets of spectra. This fragmentation map, hereafter called fragmap, provides a generalization of a mirror plot but also improves upon the visual use of the space available for plotting. This is done by employing a factorized and sorted recasting of binned mass-to-charge ratios. Here, factorized mass-to-charge ratios are put onto the x-axis in order, while the y-axis delineates the spectra. For each mass-to-charge ratio bin, intensities of the fragment in the respective spectra are displayed using a color gradient as typical in heatmaps. Fragment overlap can be assessed instantly for each fragment between spectra, while intensity agreements or disagreements are easily spotted. Importantly, the x-axis is the same for all spectra and the plot loses minimal plotting surface to white spaces as distance in mass to charge ratios between fragments is removed through factorization. Optionally, each spectrum is filtered down to its respective top-K highest intensity fragments before factorization and neutral loss computation to provide a simplified view for complex or noisy spectra. The result is a concise overview of fragmentation overlaps between spectra for quick assessments of relatedness between matched nodes. While fragmaps provide a means of quickly asserting overlaps across spectra, they perform a number of transformation steps that can obscure the raw data. Most notably, binning may aggregate different fragment peaks into the same bin, leading to loss of information and possibly misleading overlaps in the fragmentation overview. This potential bias is counteracted using interactive hover pop-ups that provide the original mass-to-charge ratio values and intensities of any fragments being aggregated into a single bin, for each bin-fragment and spectrum separately. The user can thus quickly assess the bin artefacts when inspecting promising overlaps. In addition, to provide a more direct and familiar link to the raw spectra themselves, specXpolore also provides spectrum plots and mirror plots.

### 5 Tabular Metadata Integration

Not all metadata lends itself well to graphical visualization in the spectral exploration dashboard, but may nonetheless be useful to have integrated into the tool for quick access upon node selection. SpecXplore offers a metadata table panel that can be generated for any node selection, allowing the user to inspect any additional supplied information regarding a spectrum inside the dashboard. This is where ms2query analog hits, but also any user-provided metadata can be displayed for selections of nodes. Incorporation of this table allows users to explore their data within specXplore and thus removes some of the need for switching between programs. That being said, it also allows users to inspect relevant identifiers in the metadata to quickly inspect features in other programs as well.

### 6 Spectral data pre-processing

SpecXplore makes use of pairwise similarity computations and direct spectral comparison approaches that require some spectral data pre-processing. First, spectra are normalized such that intensities are measured relative to the most intense fragment. This is necessary for some pairwise similarity matrices, and fragmap overviews. Subsequently, the spectra are filtered to only keep fragments if their relative intensity exceeds some set minimum or to limit the spectrum to the top n (default of two-hundred) fragments ranked by fragment intensity. This step is done in order to a) limit noise peaks from diluting signals that interfere with pairwise spectral similarity measure computations, and b) limit very large and noisy spectra from diluting and obscuring fragmentation overview visualizations while contributing little effective insight. Finally, any fragments outside the mass-to-charge ratio window of zero to one thousand will be removed as they are not supported by the used machine learning model and fall outside the spectral binning range. Additional processing such a minimum number of fragments per spectrum may also be applied to focus analysis on more promising and information rich spectra for analysis. It should be noted that these processing steps happen in specXplore on the basis of .mgf spectral exports. Whenever possible, pre-processing should be done in software operating closer to the raw data, such as e.g. MZMine [2], as more relevant information for filtering and quality checks will be available.

### 7 Usage and tuning of t-SNE as a fixed network layout

Finding good settings for t-SNE is a task that is difficult to automate and may even warrant a dashboard of its own to formally evaluate and understand as done in t-viSNE [4]. In our trials, we've noticed that often times very high perplexities lead to the best distance preservation for the t-SNE embedding. However, larger perplexity usually implies larger groups of nodes and hence larger overlaps between topological groups. Hence, the best perplexity settings for t-SNE require an informal trade-off in distance preservation and visual characteristics. Despite lacking formally optimal settings for the used t-SNE embedding, we explicitly chose to make the t-SNE embedding in specXplore fixed rather than modifiable on the fly. This has three reasons; computational speed, robustness, and mental map preservation. First, t-SNE embedding computations can be computationally costly for larger datasets and may require long run-times. In specXplore and dash, long run times may be easy to confuse with app crashes. In addition, t-SNE embeddings may require a number of t-SNE runs to be able to assess what embedding works well in a qualitative sense. This effectively multiplies the computational speed bottleneck of t-SNE tuning in the app. These two aspects together make tuning prohibitive in an interactive session. Second, during our trials t-SNE runs showed surprising qualitative robustness, often leading to useful embeddings regardless of precise perplexity values used. Third, the t-SNE embedding is meant to provide a global overview from which to explore the dataset. Should nodes be scattered differently, the user's 'mental map' and hence overview of the data would be lost. Given these considerations, it seemed most appropriate to defer t-SNE tuning to the jupyter notebook pre-processing step. Alternative approaches to data visualization were considered such as tmap [5, 6] or pacmap [7]. However, given the precedent of t-SNE in the field for instance MetGem [8], as well as a combination of good agreement with local topology at higher thresholds and straightforward compatibility with arbitrary similarity matrices, made t-SNE our method of choice. Alternative embedding approaches, include those approaches that avoid the computation of a full pairwise similarity matrix, may be especially useful in larger visualization contexts going beyond individual experiment visualization.

### 8 In-silico spike-in standards

SpecXplore includes the option of using in-silico spike-in reference standards in the spectral data exploration workflow. This is done via adding additional spectra and corresponding metadata and assigning the add-on features as features to be-highlighted in the t-SNE overview. This option combines local library search and molecular networking style visualization in a single exploratory visualization approach. Naturally, such standards can be from any online repository and instrument type, but can also be from in-house libraries affording greater spectral similarity through similar or equal instrumental setups being used in standard and experimental spectra acquisition. By making reference standards stand out visually using higher opacity and darker grays, chemical spaces of interest can be quickly identified in the global overview.

### 9 Color as a medium for prioritization and visual emphasis

In general, color is one of the most effective visual channels available to us; both for categorical and continuous data. In specXplore, we use color as a tool for emphasis and guiding attention. For instance, spike-in standards are highlighted in darker gray colors and increased opacity compared to experimental nodes. Any selected nodes are encircled with magenta highlighting. In egonets, we make use of the Viridis color scheme discretized to n colors, where n is the number of hops allowed, to indicate distance from the egonet. Here, color is further reinforced by opacity and line thickness changes as a function of hop distance. In addition, network views for selections of nodes emphasize the selected nodes and their interconnections in magenta while connections out of the selected set are visualized in black. Chemical classes or k-medoids groupings can further be highlighted in color within the t-SNE overview graph. While color provides a powerful medium for emphasis, care must be taken to avoid "color saturation", as humans are generally only able to meaningfully differentiate up to eight to twelve unique colors [9]. Additionally, color-blind-friendly color-maps should be used as much as possible [10, 11]. In specXplore, we make use of the *ColorBrewer* [12] Dark2 to provide an appropriate color palette for highlighting up to eight different node classes in the overview t-SNE graph. In addition, to make our augmented heatmap representation more useful to colorblind users, we incorporated a toggle for grayscale similarity depiction with magenta markers, thus avoiding combinations of blue, red, and green at the same time.

### 10 Chemical Class Visualization

Chemical ontologies provide a scientific foundation for data subdivision [13, 14]. However, ontologies require known chemical structures which are generally not available for experimental mass spectral data explored using molecular networking. Different means of providing chemical information for unknown spectra exist such as Canopus for chemical classification [15, 16] or ms2lda for chemical motif detection [17]. In specXplore, the exact use of predicted ontology is left to the user and easy to integrate via feature identifier matching. The ms2query best-scoring analog classifications may be used to get a rough feeling for chemical class via putative analog class distribution in the overview graph. A current limitation in specXplore is that, at each ontology level, each feature may only be member of a single class. In practice, this means that any multiple classifications are amalgamated together into a single entry for each feature. This may lead to some loss of class select-ability for highlighting.

### 11 Effective visual analysis using scented widgets

Effective visual analysis depends on both the selected visual encoding and the means for interactively traversing said encoding. A host of possible interaction techniques present themselves to make this navigation as effective as possible [18, 19]. For specXplore, localized on-demand selections and interactive filtering of edges between nodes is used for allowing users to explore local graph topology without excessive visual clutter. Determining what threshold to use in graph visualization is a non-trivial task [20]. Inspired by Willet et al.'s *Scented Widgets* [21], specXplore provides a visual summary of the edge weight distribution to the input slider in order to provide insights into global impacts of thresholds on edge numbers. For more local impact evaluations, node degrees can be visually highlighted using a color gradient from the minimum node size to the maximum node size in the t-SNE overview. Both of these options, as well as the augmap-based matrix representation, provide insights into the topological impacts of thresholds and thus allow exploration in an informed manner rather than random trial and error. Of course, metabolomics domain considerations will ultimately determine the acceptance or rejection of edges.

### 12 Augmented Matrix Views

We make use of a multilevel heatmap representation of the pairwise similarity matrix to facilitate quantitative insight generation into the actual ms2deepscore pairwise similarity matrix, but also to allow comparison between scores from the ms2deepscore anchor point. Different similarity measures for the same features can be viewed together as a multi-layer graph, i.e. a set of nodes connected by different "layers" of edges (in this case) defined by each similarity measure [22]. Visualizing such graphs is non-trivial, and a number of visualization techniques have been put forth [23]. In specXplore, our graph comparison aims to imply that we are interested in visualizing all the layers of the multi-graph simultaneously. Given the visual clutter that rendering all such edges would introduce, a conventional network representation would be rendered nigh unreadable. Similarly, radial embeddings [24] as well as so-called *Hive Plots* [25] were considered, but ultimately deemed inappropriate. Instead, we opt for a matrix representation, as they have been shown to be more effective and preferred (in all but path-tracing tasks [26]) when applied to large and/or dense graphs [27, 28]. Importantly, when using the matrix representation in the form of a heatmap, quantitative information can be color encoded for one of the measures, with additional markers giving qualitative adjacency insights into the other metrics. Such marker-augmented heatmaps are uncommon, though there is precedent in other fields (e.g. figure 5 in [29]).

### 13 Bibliography


[1] Ian Oesterle, Manuel Pristner, Sabrina Berger, Mingxun Wang, Vinicius Verri Hernandes, Annette Rompel, and Benedikt Warth. Exposomic biomonitoring of polyphenols by non-targeted analysis and suspect screening. *Analytical Chemistry*, 95(28):10686–10694, 2023.

[2] Robin Schmid, Steffen Heuckeroth, Ansgar Korf, Aleksandr Smirnov, Owen Myers, Thomas S. Dyrlund, Roman Bushuiev, Kevin J. Murray, Nils Hoffmann, Miaoshan Lu, Abinesh Sarvepalli, Zheng Zhang, Markus Fleischauer, Kai Dührkop, Mark Wesner, Shawn J. Hoogstra, Edward Rudt, Olena Mokshyna, Corinna Brungs, Kirill Ponomarov, Lana Mutabdžija, Tito Damiani, Chris J. Pudney, Mark Earll, Patrick O. Helmer, Timothy R. Fallon, Tobias Schulze, Albert Rivas-Ubach, Aivett Bilbao, Henning Richter, Louis-Félix Nothias, Mingxun Wang, Matej Orešič, Jing-Ke Weng, Sebastian Böcker, Astrid Jeibmann, Heiko Hayen, Uwe Karst, Pieter C. Dorrestein, Daniel Petras, Xiuxia Du, and Tomáš Pluskal. Integrative analysis of multimodal mass spectrometry data in MZmine 3. *Nature Biotechnology*, 41(4):447–449, March 2023. doi: 10.1038/s41587-023-01690-2. URL https://doi.org/10.1038/s41587-023-01690-2.

[3] Alexander Lex, Nils Gehlenborg, Hendrik Strobelt, Romain Vuillemot, and Hanspeter Pfister. UpSet: Visualization of Intersecting Sets. *IEEE Transactions on Visualization and Computer Graphics*, 20(12): 1983–1992, December 2014. ISSN 1941-0506. doi: 10.1109/TVCG.2014.2346248. Conference Name: IEEE Transactions on Visualization and Computer Graphics.

[4] Angelos Chatzimparmpas, Rafael M. Martins, and Andreas Kerren. t-visne: Interactive assessment and interpretation of t-sne projections. *IEEE Transactions on Visualization and Computer Graphics*, 26(8): 2696–2714, 2020. doi: 10.1109/TVCG.2020.2986996.

[5] Daniel Probst and Jean-Louis Reymond. Visualization of very large high-dimensional data sets as minimum spanning trees. *Journal of Cheminformatics*, 12(1):12, Feb 2020. ISSN 1758-2946. doi: 10.1186/s13321-020-0416-x. URL https://doi.org/10.1186/s13321-020-0416-x.

[6] Pierre-Marie Allard, Arnaud Gaudry, Luis-Manuel Quirós-Guerrero, Adriano Rutz, Miwa Dounoue-Kubo, Tom W N Walker, Emmanuel Defossez, Christophe Long, Antonio Grondin, Bruno David, and Jean-Luc Wolfender. Open and reusable annotated mass spectrometry dataset of a chemodiverse collection of 1,600 plant extracts. *GigaScience*, 12, 01 2023. ISSN 2047-217X. doi: 10.1093/gigascience/giac124. URL https://doi.org/10.1093/gigascience/giac124. giac124.

[7] Yingfan Wang, Haiyang Huang, Cynthia Rudin, and Yaron Shaposhnik. Understanding how dimension reduction tools work: an empirical approach to deciphering t-sne, umap, trimap, and pacmap for data visualization. *The Journal of Machine Learning Research*, 22(1):9129–9201, 2021.

[8] Florent Olivon, Nicolas Elie, Gwendal Grelier, Fanny Roussi, Marc Litaudon, and David Touboul. Metgem software for the generation of molecular networks based on the t-sne algorithm. *Analytical Chemistry*, 90 (23):13900–13908, 2018. doi: 10.1021/acs.analchem.8b03099. URL https://doi.org/10.1021/acs.analchem.8b03099. PMID: 30335965.

[9] Tamara Munzner. *Visualization Analysis and Design*. A K Peters/CRC Press, New York, October 2014. ISBN 978-0-429-08890-2. doi: 10.1201/b17511.

[10] Luke Jefferson and Richard Harvey. Accommodating color blind computer users. In *Proceedings of the 8th international ACM SIGACCESS conference on Computers and accessibility*, Assets '06, pages 40–47, New York, NY, USA, October 2006. Association for Computing Machinery. ISBN 978-1-59593-290-7. doi: 10.1145/1168987.1168996. URL https://doi.org/10.1145/1168987.1168996.

[11] Frank Elavsky, Cynthia Bennett, and Dominik Moritz. How accessible is my visualization? Evaluating visualization accessibility with Chartability. *Computer Graphics Forum*, 41(3):57–70, 2022. ISSN 1467-8659. doi: 10.1111/cgf.14522. URL https://onlinelibrary.wiley.com/doi/abs/10.1111/cgf.14522. _eprint: https://onlinelibrary.wiley.com/doi/pdf/10.1111/cgf.14522.

[12] Mark Harrower and Cynthia A. Brewer. ColorBrewer.org: An Online Tool for Selecting Colour Schemes for Maps. *The Cartographic Journal*, 40(1):27–37, June 2003. ISSN 0008-7041. doi: 10.1179/000870403235002042. URL https://www.tandfonline.com/doi/abs/10.1179/000870403235002042. Publisher: Taylor & Francis _eprint: https://www.tandfonline.com/doi/pdf/10.1179/000870403235002042.

[13] Yannick Djoumbou Feunang, Roman Eisner, Craig Knox, Leonid Chepelev, Janna Hastings, Gareth Owen, Eoin Fahy, Christoph Steinbeck, Shankar Subramanian, Evan Bolton, Russell Greiner, and David S. Wishart. Classyfire: automated chemical classification with a comprehensive, computable taxonomy. *Journal of Cheminformatics*, 8(1):61, Nov 2016. ISSN 1758-2946. doi: 10.1186/s13321-016-0174-y. URL https://doi.org/10.1186/s13321-016-0174-y.

[14] Hyun Woo Kim, Mingxun Wang, Christopher A. Leber, Louis-Félix Nothias, Raphael Reher, Kyo Bin Kang, Justin J. J. van der Hooft, Pieter C. Dorrestein, William H. Gerwick, and Garrison W. Cottrell. Npclassifier: A deep neural network-based structural classification tool for natural products. *Journal of Natural Products*, 84(11):2795–2807, 2021. doi: 10.1021/acs.jnatprod.1c00399. URL https://doi.org/10.1021/acs.jnatprod.1c00399. PMID: 34662515.

[15] Kai Dührkop, Markus Fleischauer, Marcus Ludwig, Alexander A. Aksenov, Alexey V. Melnik, Marvin Meusel, Pieter C. Dorrestein, Juho Rousu, and Sebastian Böcker. Sirius 4: a rapid tool for turning tandem mass spectra into metabolite structure information. *Nature Methods*, 16(4):299–302, Apr 2019. ISSN 1548-7105. doi: 10.1038/s41592-019-0344-8. URL https://doi.org/10.1038/s41592-019-0344-8.

[16] Kai Dührkop, Louis-Félix Nothias, Markus Fleischauer, Raphael Reher, Marcus Ludwig, Martin A Hoffmann, Daniel Petras, William H Gerwick, Juho Rousu, Pieter C Dorrestein, and Sebastian Böcker. Systematic classification of unknown metabolites using high-resolution fragmentation mass spectra. *Nat Biotechnol*, 39(4):462–471, November 2020.

[17] Justin Johan Jozias van der Hooft, Joe Wandy, Michael P. Barrett, Karl E. V. Burgess, and Simon Rogers. Topic modeling for untargeted substructure exploration in metabolomics. *Proceedings of the National Academy of Sciences*, 113(48):13738–13743, 2016. doi: 10.1073/pnas.1608041113. URL https://www.pnas.org/doi/abs/10.1073/pnas.1608041113.

[18] Ji Soo Yi, Youn ah Kang, John Stasko, and J.A. Jacko. Toward a Deeper Understanding of the Role of Interaction in Information Visualization. *IEEE Transactions on Visualization and Computer Graphics*, 13 (6):1224–1231, November 2007. ISSN 1941-0506. doi: 10/fmrs6r. Conference Name: IEEE Transactions on Visualization and Computer Graphics.

[19] Ana Figueiras. Towards the Understanding of Interaction in Information Visualization. In *2015 19th International Conference on Information Visualisation*, pages 140–147, July 2015. doi: 10.1109/iV.2015.34. ISSN: 2375-0138.

[20] Jinwook Seo and B. Shneiderman. A Rank-by-Feature Framework for Unsupervised Multidimensional Data Exploration Using Low Dimensional Projections. In *IEEE Symposium on Information Visualization*, pages 65–72, October 2004. doi: 10.1109/INFVIS.2004.3. ISSN: 1522-404X.

[21] Wesley Willett, Jeffrey Heer, and Maneesh Agrawala. Scented Widgets: Improving Navigation Cues with Embedded Visualizations. *IEEE Transactions on Visualization and Computer Graphics*, 13(6):1129–1136, November 2007. ISSN 1941-0506. doi: 10.1109/TVCG.2007.70589. Conference Name: IEEE Transactions on Visualization and Computer Graphics.

[22] Mikko Kivelä, Alex Arenas, Marc Barthelemy, James P. Gleeson, Yamir Moreno, and Mason A. Porter. Multilayer networks. *Journal of Complex Networks*, 2(3):203–271, September 2014. ISSN 2051-1310. doi: 10.1093/comnet/cnu016. URL https://doi.org/10.1093/comnet/cnu016.

[23] F. McGee, M. Ghoniem, G. Melançon, B. Otjacques, and B. Pinaud. The State of the Art in Multilayer Network Visualization. *Computer Graphics Forum*, 38(6):125–149, 2019. ISSN 1467-8659. doi: 10.1111/cgf.13610. URL https://onlinelibrary.wiley.com/doi/abs/10.1111/cgf.13610. _eprint: https://onlinelibrary.wiley.com/doi/pdf/10.1111/cgf.13610.

[24] Martin Krzywinski, Jacqueline Schein, İnanç Birol, Joseph Connors, Randy Gascoyne, Doug Horsman, Steven J. Jones, and Marco A. Marra. Circos: An information aesthetic for comparative genomics. *Genome Research*, 19(9):1639–1645, September 2009. ISSN 1088-9051, 1549-5469. doi: 10.1101/gr.092759.109. URL https://genome.cshlp.org/content/19/9/1639. Company: Cold Spring Harbor Laboratory Press Distributor: Cold Spring Harbor Laboratory Press Institution: Cold Spring Harbor Laboratory Press Label: Cold Spring Harbor Laboratory Press Publisher: Cold Spring Harbor Lab.

[25] Martin Krzywinski, Inanc Birol, Steven JM Jones, and Marco A Marra. Hive plots—rational approach to visualizing networks. *Briefings in Bioinformatics*, 13(5):627–644, September 2012. ISSN 1467-5463. doi: 10.1093/bib/bbr069. URL https://doi.org/10.1093/bib/bbr069.

[26] Mohammad Ghoniem, Jean-Daniel Fekete, and Philippe Castagliola. On the Readability of Graphs Using Node-Link and Matrix-Based Representations: A Controlled Experiment and Statistical Analysis. *Information Visualization*, 4(2):114–135, June 2005. ISSN 1473-8716, 1473-8724. doi: 10.1057/palgrave.ivs.9500092. URL http://journals.sagepub.com/doi/10.1057/palgrave.ivs.9500092.

[27] René Keller, Claudia M. Eckert, and P. John Clarkson. Matrices or Node-Link Diagrams: Which Visual Representation is Better for Visualising Connectivity Models? *Information Visualization*, 5(1):62–76, March 2006. ISSN 1473-8716, 1473-8724. doi: 10.1057/palgrave.ivs.9500116. URL http://journals.sagepub.com/doi/10.1057/palgrave.ivs.9500116.

[28] M. Ghoniem, J.-D. Fekete, and P. Castagliola. A Comparison of the Readability of Graphs Using Node-Link and Matrix-Based Representations. In *IEEE Symposium on Information Visualization*, pages 17–24, October 2004. doi: 10.1109/INFVIS.2004.1. ISSN: 1522-404X.

[29] Bumhee Park, Dae-Shik Kim, and Hae-Jeong Park. Graph independent component analysis reveals repertoires of intrinsic network components in the human brain. *PLOS ONE*, 9(1):1–10, 01 2014. doi: 10.1371/journal.pone.0082873. URL https://doi.org/10.1371/journal.pone.0082873.
